# Best practices for improving alignment and variant calling on human sex chromosomes

**DOI:** 10.1101/2025.04.29.651297

**Authors:** Angela M. Taravella Oill, Seema B. Plaisier, Tanya N. Phung, Melissa A. Wilson

**Affiliations:** Center for Evolution and Medicine, Arizona State University, Tempe, AZ 85281 USA; School of Life Sciences, Arizona State University, Tempe, AZ 85281 USA; Comparative Genomics and Reproductive Health Section, Center for Genomics and Data Science Research, National Human Genome Research Institute, National Institutes of Health, Bethesda, MD, 20852 USA

**Keywords:** Alignment, variant calling, variant filtering, sex chromosomes, X chromosome, Y chromosome

## Abstract

Sex chromosome complement is the largest karyotypic variation observed in humans. X and Y chromosomes were once a pair of homologous autosomes. Although chromosome X and Y differentiated from one another, they still share high levels of sequence similarity in some regions, like the pseudoautosomal regions (PARs) and the X-transposed region (XTR). The sex chromosomes violate some assumptions of autosomal pairs, but are not always processed separately in genomics analyses. Here, we undertook a simulation study to assess the effects of standard autosomal versus sex chromosome complement-informed alignment, variant calling, and variant filtering strategies on variants detected on the human sex chromosomes. We find that aligning samples to a reference genome informed by the sex chromosome complement of the sample increases the number of true positives called in the PARs, and, in XX-samples only, also the XTR. In contrast, in XY-samples, masking the XTR during alignment results in a ten-fold higher rate of false positives. We further find that haploid calling on the sex chromosomes in XY-samples reduces the number of false positives compared to diploid calling, but does not decrease the number of false negatives. Improving the accuracy of variant calling results in detection of variants that could be relevant to studies of health and disease, including variants we recovered in genes implicated in cardiomyopathy, immunodeficiency, and Alzheirmer’s disease, among others. We recommend future genomic analyses implement the following best practices for detecting variants: aligning samples to versions of the human reference genome informed by the sex chromosome complement of the sample and using accurate ploidy parameters when calling variants.

## Introduction

Despite significant sex differences in nearly every disease including cancer (Lopes-Ramos, Quackenbush, and DeMeo 2020), cardiovascular disease (Regitz-Zagrosek and Kararigas 2017), aging and immune function (Yanicke and Ucar 2023), and Alzhimer’s disease (Oveisgharan et al. 2018; Liesinger et al. 2018), the X and Y chromosomes are routinely excluded or incompletely included. The sex chromosomes, X and Y, in particular, contain distinct information from the autosomes (non-sex chromosomes) (Musharoff et al. 2019; Wilson Sayres, Lohmueller, and Nielsen 2014). There are efforts to develop better approaches to incorporating sex chromosomes into human disease genetics (Khramtsova et al. 2023; Khramtsova, Davis, and Stranger 2019), but a root challenge in finding associations between sex chromosome variants and disease is first being able to identify the variants. Alignment, variant calling, and filtering are essential steps in generating an accurate variant call set for genomic analysis of human disease. Single nucleotide variants are one type of genetic variant used in genomics analyses that can provide important insights into normal human health and development, the genetic basis of complex diseases, or can be used in population genetics analysis (Jackson et al. 2018; Timpson et al. 2018; Hindorff, Gillanders, and Manolio 2011; Shastry 2002). However, the sex chromosomes present unique challenges in alignment, variant calling, and filtering (Webster et al. 2019; Pinto et al. 2023a; Olney et al. 2020; Khramtsova et al. 2023).

Therian mammalian X and Y chromosomes share an evolutionary origin as they were once a pair of homologous autosomes recombining across their entire lengths (Ohno 2013). Around 180-120 million years ago sex determining genes were acquired and recombination suppression led to the eventual differentiation of these chromosomes (Livernois, Graves, and Waters 2012; Rens et al. 2007; Charlesworth 1991). Despite this differentiation, the X and Y chromosomes are still homologous and recombine at each end of their chromosomes - these are known as the pseudoautosomal regions (PARs) (Ross et al. 2005). There are also other regions of X and Y with high sequence similarity like the X-transposed region, which share about 98.78% sequence similarity (Veerappa, Padakannaya, and Ramachandra 2013). This unique evolutionary history and regions of sequence similarity pose methodological and analytical challenges for genomic analysis of the sex chromosomes (Khramtsova, Davis, and Stranger 2019). Sequence homology affects read mapping and subsequent variant calling on the sex chromosomes (Webster et al. 2019). However, previous work did not explicitly assess whether variant calling was improved with this approach as there was no truth set of variants to compare to and variant calling and filtering approaches were not investigated.

Best practices and workflows for alignment (Li and Homer 2010), variant calling (Koboldt 2020; Kumaran, Subramanian, and Devarajan 2019; Pirooznia et al. 2014; Liu et al. 2013; DePristo et al. 2011) and filtering (Jia et al. 2012; Reumers et al. 2011; DePristo et al. 2011) have been produced for the autosomes, but not the sex chromosomes. Here we assessed whether aligning samples to reference genomes informed by the sex chromosome complement of the sample improves variant calling compared to aligning samples to a default reference genome. In addition to sequence homology on the sex chromosomes, the X chromosome is diploid in XX-samples while the X and Y chromosomes are largely haploid in XY-samples. Thus, we further investigated whether, and how, diploid calling and diploid-based filtering threshold on haploid chromosomes impact variant calling.

To assess and outline the best practices for mapping, calling and filtering variants on the human sex chromosomes we simulated genome-wide sequence reads for 60 female (46, XX) and 60 male (46, XY) samples to 20x coverage and compare how taking a standard approach for alignment and variant calling affects variant calling on the sex chromosomes compared to a sex-chromosome complement aware approach (**Figure 1**). We report here that aligning samples to a reference genome informed on the sex chromosome complement of the sample, using biological relevant ploidy for calling variants, and setting filtering thresholds informed by ploidy of the chromosome improves variant calling on the sex chromosomes. We find that aligning samples to a reference genome informed by the sex chromosome complement of the sample increases the number of true positives called in the PARs, and additionally, in XX-samples only, the XTR. We find that haploid calling on X and Y in XY-samples reduces the number of false positives compared to diploid calling but does not substantially decrease the number of false negatives. Additionally, we found that implementing diploid-based thresholds for total depth (DP) and allele number (AN) on the sex chromosomes in XY-samples results in a higher loss of true positives compared to when implementing haploid-based thresholds. Finally, we tested a new sex-aware approach for attempting to improve variant calling in XY-samples in the XTR - a region that shows high sequence similarity but not complete identity between X and Y. We show that masking one copy of the XTR when aligning XY-samples results in a ten-fold higher rate of false positives in the XTR and so should not be implemented. Based on these results, our recommendation is to align samples to versions of the human reference genome informed on the sex chromosome complement of the sample and to use karyotypically accurate ploidy parameters when calling variants and setting filtering thresholds.

**Figure 1.**
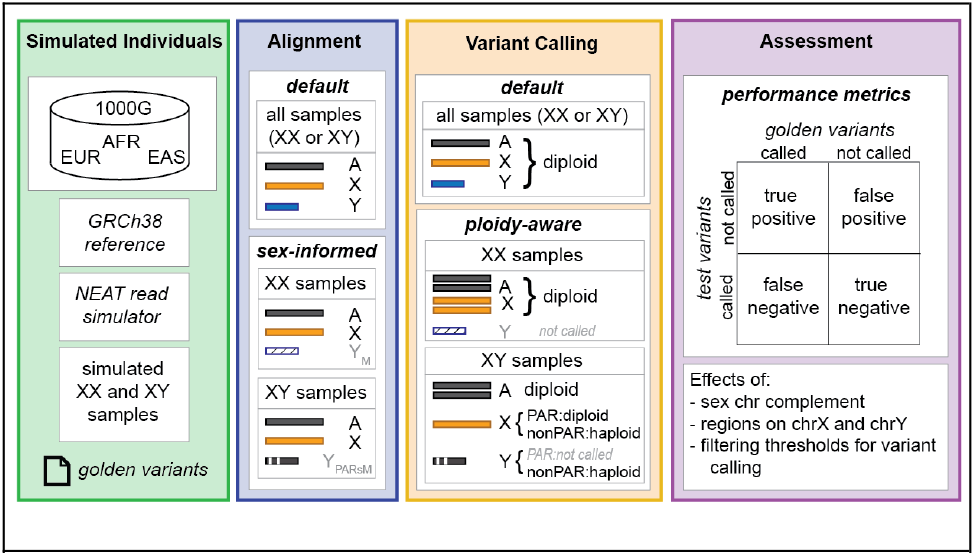
Schematic overview of sex chromosome variant simulations, variant calling, and assessment. Here we summarize the simulations and assessments conducted here. We begin (Simulated individuals) by generating a set of golden alignment (BAM) and variant (VCF) files containing the “true” simulated read alignments and variants. We the (Alignment) implemented three alignment approaches, including a default autosomal approach, and two alternative sex chromosome complement informed approaches, based on the presence or absence of a Y chromosome. Next (Variant Calling) we implemented default autosomal ploidy assumptions across all chromosomes in all samples, versus using chromosome-informed ploidy for diploid/haploid regions of the X and Y in XY-samples. Finally (Assessment), we compared the test variants (called and not called) versus the golden variants (called and not called) to assess the effects of each decision point in alignment and variant calling.

## Materials and Methods

### Genomic features on X and Y

Pseudoautosomal regions (PARs), X-transposed regions (XTR) and ampliconic regions were used for both analysis and visualizations. PAR coordinates on X and Y were obtained from Ensembl GRCh38 PAR definitions (Aken et al. 2017) (**Table S3**). XTR and ampliconic coordinates on the Y chromosome were obtained from Poznik et al. (Poznik et al. 2013), XTR coordinates on the X chromosome were obtained from Webster et al. (Webster et al. 2019), and ampliconic coordinates on the X chromosome were obtained from Cotter et al. (Cotter, Brotman, and Wilson Sayres 2016) based on Muller et al. (Mueller et al. 2013) in hg19. XTR and ampliconic coordinates were lifted over from hg19 to GRCh38 using UCSC LiftOver tool (Kent et al. 2002) (**Table S3**).

The XTR on the X and Y chromosomes and ampliconic regions on the Y chromosome successfully lifted over. On the X chromosome, 4 out of 14 ampliconic regions failed to lift over when using the default LiftOver options (ampliconic region 2, 3, 5, and 7 in **Table S3**). For the four that failed lift over, the error message reported was “split in new”, so we ran LiftOver again for these four, specifying “Allow multiple output regions” option. This resulted in 3 of the 4 amplicons successfully lifting over (**Table S3**). Ampliconic region 7 was split between both chromosome X and chromosome 1 so we only analyzed the region that lifted over on chromosome X. Ampliconic region 3 still failed lift over so we did not include it in our visualizations and analyses. Annotations are included on the Github repository for this project: https://github.com/SexChrLab/VariantCalling

### Simulating sequence reads

We simulated paired-end sequence reads with a read length of 150 base pairs (bp) using NExt-generation sequencing Analysis Toolkit (NEAT) software version 3.0 (Stephens et al. 2016). Variants were inserted from 20 males and 20 females each, from European, African, and East Asian populations (a total of 120 individuals) from high coverage variant calls from 1000 Genomes (Byrska-Bishop et al. 2021) using the “-v” option in NEAT (**Table S2**). NEAT requires a reference genome to sample reads from, so we used the same reference genome that was used in the generation of the 1000 Genomes high coverage GRCh38 VCFs (ftp://ftp.1000genomes.ebi.ac.uk/vol1/ftp/technical/reference/GRCh38_reference_genome/GRCh38_full_analysis_set_plus_decoy_hla.fa). Additionally, we used the default sequencing error model derived from illumina reads provided in the NEAT software. For the autosomes, mitochondrial DNA (mtDNA), and X non-PARs (females only) coverage was set to 20x. For the X and Y non-PARs in males, coverage was set to 10x each as both are haploid in a 46, XY genome and expected to have half the sequencing coverage of a diploid autosome. PAR sequences were simulated as diploid on the X chromosome for both males and females because the PAR variants in 1000 genomes were called only on the X chromosome. NEAT produces simulated FASTQs, along with a golden VCF with the set of true positive variants and a golden BAM with a golden set of aligned reads.

### Trimming and alignment

Using bbduk - a tool within bbmap version 38.92 - (Bushnell 2014), bases with a Phred score less than 20 were trimmed on both the left and right side of reads, reads shorter than 75bp were discarded, and reads with average Phred quality below 20 were removed (bbduk trimming parameters used: “qtrim=rl trimq=20 minlen=75 maq=20”). Reads were mapped to one of three versions of a reference genome: a version of 1000 Genomes GRCh38 reference genome where the both X and Y chromosomes were unmasked (default reference) and versions informed by the sex chromosome complement (SCC reference) of the sample (either with or without evidence of a Y chromosome). The GRCh38 reference genome provided by 1000 Genomes has the Y chromosome PARs already hard masked, so to make a default version of this reference genome - one where the Y PARs are unmasked - we downloaded the original reference genome from GenBank (NCBI assembly accession: GCA_000001405.15) and replaced the Y chromosome with the unmasked Y sequence. The sex chromosome complement reference genome for females (XX) has the Y chromosome hard masked out, while the sex chromosome complement reference genome for males (XY) has both of the PARs on the Y chromosome hard masked (Webster et al. 2019). For simulated male samples only, we additionally aligned reads to the XY-SCC version of 1000 Genomes GRCh38 reference genome with the homologous Y XTR sequence hard masked out (SCC + Y-XTR masked reference). The Y chromosome XTR was hard masked using bedtools v2.30.0 maskfasta (Quinlan and Hall 2010). All alignments were performed using Burrows-Wheeler Aligner software (https://doi.org/10.1093/bioinformatics/btp324).

### Variant calling and filtering

Variant calling was performed using GATK’s v4.2.1.0 HaplotypeCaller tool, joint genotyping was performed using GenotypeGVCFs tool, and filtering was performed using SelectVariants and VariantFiltration tools (McKenna et al. 2010). In the standard approach we processed chromosome X and Y in the same way as the autosomes: variants were called in diploid mode, joint genotyped together, and filtered using GATKs hard filtering autosomal threshold recommendations (filtering parameters: -filter “QD < 2.0” -filter “QUAL < 30.0” -filter “SOR > 3.0” -filter “FS > 60.0” -filter “MQ < 40.0” -filter “MQRankSum < −12.5” -filter “ReadPosRankSum < −8.0”). For males we also implemented a haploid approach where everything remained the same as the standard approach except the non-PAR X and Y chromosomes were called in haploid mode. In addition to GATK’s recommended thresholds for hard filtering, we also tested different thresholds for a subset of the hard filters (QualByDepth) and tested additional filters (allele number (AN) and total depth (DP)) on our ability to accurately call variants in the simulated data. For QualByDepth we tested the following thresholds: 1, 1.5, 2, 12, 16, 20, 28, for allele number we tested the following thresholds: 20, 15, 10, 5, 4, 3, 2, 1, and for total depth we implemented diploid- and haploid-based thresholds. The diploid-based threshold was based on the mean DP across the autosomes, retaining sites whose DP fell between 50% and 150% mean DP. The haploid-based threshold was based on the mean DP across X and Y (for males only), retaining sites whose DP fell between 50% and 150% mean DP.

### Calculations and comparisons of performance metrics

We calculated true positives (TP), false positives (FP), and false negatives (FN) for each individual, using a custom python script. Using both the simulated VCFs output from NEAT that have the ground truth variants for each individual and the called VCFs from the different approaches, a TP was defined as a SNP that was both called and simulated and where the reference and alternative allele matched between the golden and called VCFs, a FP was defined as a SNP called but not simulated, and a FN was defined as a SNP that was simulated but not called.

To visually assess performance metrics, we counted FNs, FPs, and TPs across X and Y chromosomes across 50kb windows using bedtools map v2.30.0 (Quinlan and Hall 2010). All plotting of performance metrics were performed in R (Computing and Others 2013).

## Results

### There are more true positive on chromosome X when aligning samples to a sex chromosome complement reference genome compared to a default reference genome

Compared to the default alignment, we find that variant calling is improved on the X chromosome when aligning samples to a reference genome informed on the sex chromosome complement of the sample. We observe more true positives in PARs for both XX and XY samples and XTR in XX samples only when aligning to a reference genome informed on the sex chromosome complement of the sample compared to default alignment (**Figure 2A and 2B**). There are on average 4909 and 4980 more true positives in the PARs in simulated females and males, respectively, when aligning samples to a sex chromosome complement reference genome compared to a default reference (**Table 1**). In the XTR on the X chromosome, there are on average 748 more true positives in females while for males, there is no difference in the number of true positives between alignment strategies (**Table 1**). For the non-PAR region of the Y chromosome, there are no differences in the number of true positives between alignment strategies (**Figure 2C; Table 1**). We replicate this analysis for multiple subsets of data, including in 10 or 20 male samples, 10 or 20 female samples, and when joint genotyping across the males and females together (**Table S1**). We further report the individual number of true positive variants, false positive variants, and false negative variants relative to the total number of simulated variants for the sex chromosome complement approach in each individual sample across regions of the sex chromosomes (**Table S2**).

**Table 1.** Aligning reads to sex chromosome complement reference genome improves variant calling on the X chromosome compared to aligning to a default reference genome. Average number of true positives, false positives, and false negatives on the X chromosome across 10 simulated female (XX) samples and 10 simulated male (XY) samples and across the Y chromosome in 10 simulated male (XY) samples using a default and sex chromosome complement alignment approaches. Chromosome X was split into PARs, XTR, and non-PARs without XTR and chromosome Y was split into XTR, and non-PARs without XTR.

| Chr | Region | Called in | Average # TP |  | Average # FP |  | Average # FN |  |
| --- | --- | --- | --- | --- | --- | --- | --- | --- |
|  |  |  | Default | SCC | Default | SCC | Default | SCC |
| X | PARs | Females (XX) | 0 | 4909 | 0 | 2 | 5763 | 853 |
|  |  | Males (XY) | 0 | 4980 | 0 | 3 | 5852 | 872 |
|  | non PARs (minus XTR) | Females (XX) | 97994 | 98007 | 127 | 127 | 5148 | 5135 |
|  |  | Males (XY) | 66780 | 66780 | 17 | 16 | 4217 | 4217 |
|  | XTR | Females (XX) | 3767 | 4515 | 3 | 4 | 841 | 93 |
|  |  | Males (XY) | 2171 | 2171 | 1 | 1 | 756 | 756 |
| Y | non PARs (minus XTR) | Males (XY) | 2026 | 2026 | 2 | 2 | 857 | 857 |
|  | XTR | Males (XY) | 103 | 103 | 1 | 1 | 36 | 36 |

**Figure 2.**
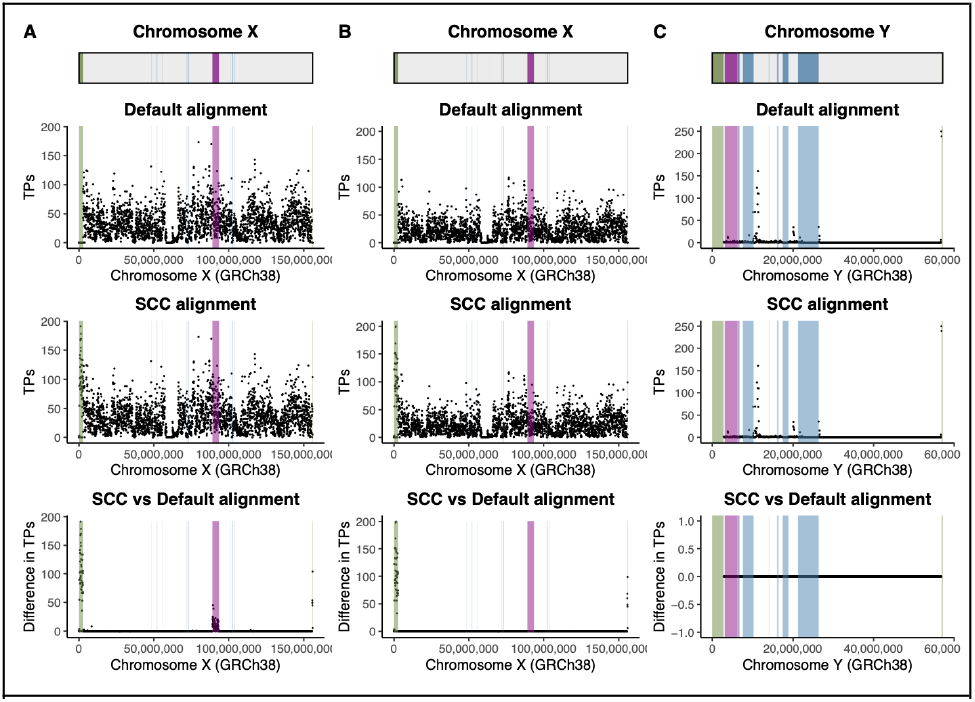
Aligning reads to a reference genome informed on the sex chromosome complement of the sample increases the number of true positives on the X chromosome compared to aligning to a default reference genome. Scatter plots showing true positives (TP) when aligning samples to default and sex chromosome complement reference genomes, and the difference in the number of true positives across chromosome X averaged across A) 10 simulated female samples and B) 10 simulated XY samples. Scatter plots were also generated for true positives when aligning samples to default and sex chromosome complement reference genomes, and the difference in the number of true positives across chromosome Y averaged across C) 10 simulated male samples. Total number of true positives were summed in 50 kb windows across the chromosomes and averaged across samples. For the difference plots (bottom figures in A-C), a negative value means there are less true positives with sex chromosome complement reference alignment compared to default reference alignment, while a positive value means there are more true positives with sex chromosome complement reference alignment compared to default reference alignment. Genomic features are annotated on the plots: green represents the PARs, purple represents XTR, and blue represents ampliconic regions. Chromosomes X and Y (non PARs) in males were called haploid, PARs were called diploid, and chromosome X was called diploid in females.

### Reduced variant calling accuracy when masking the XTR on the Y chromosome

Contrary to our expectations, when masking the X-transposed region (XTR) on the Y chromosome, accuracy declines substantially when calling variation in the XTR compared to when retaining both X-linked and Y-linked regions of the XTR. We find that there are over 10 times more variants called in the XTR than were simulated (**Figure S1**). A total of 41,236 SNPs were joint called in the XTR across the 10 male samples when aligning samples to the sex chromosome complement reference genome where the XTR on the Y chromosome was hard masked, while 6,602 and 583 SNPs were joint called in the XTR on the X and Y chromosome in male samples, respectively, when aligning sample to the sex chromosome complement reference genome without masking XTR on the Y chromosome. We also calculated true positives, false positives, and false negatives for each sample in the XTR with this alignment approach using just the X chromosome golden simulated VCFs. We find that masking the XTR on the Y chromosome resulted in an average of 33,344 more false positives compared to the sex chromosome complement approach (**Table 2; Tables S3**). This substantial increase was only observed in the XTR (**Table 2; Tables S3**).

**Table 2.** Masking XTR in the reference genome increases the number of false positives in XTR on the X chromosome while not impacting variant calling on the rest of the sex chromosomes. Mean number of true positives (TP), false positives (FP), and false negatives (FN) for the XTR on the X chromosome, X chromosome non-PAR without including the XTR and Y chromosome non-PAR without including the XTR for 10 simulated males (XY). Data were aligned in two different ways, 1) where the PARs on the Y chromosome are hard masked (referred to as Y PARs masked in the table - this is the sex chromosome complement alignment approach for XY individuals) and 2) where both Y PARs and the XTR sequence on the Y chromosome were hard masked (referred to as Y PARs + XTR masked in the table). We only assess X chromosome golden simulated variants when calculating performance metrics of XTR since variants were called in this region only on the X chromosome with the Y PARs + XTR masking approach.

| Region | Mean # TPs |  | Mean # FPs |  | Mean # FNs |  |
| --- | --- | --- | --- | --- | --- | --- |
|  | Y PARs masked | Y PARs + XTR masked | Y PARs masked | Y PARs + XTR masked | Y PARs masked | Y PARs + XTR masked |
| X<sub>XTR</sub> | 2171 | 2801 | 1 | 33345 | 756 | 98 |
| X<sub>non-PARs</sub><br>minus XTR | 66780 | 66779 | 16 | 17 | 4217 | 4218 |
| Y<sub>non-PARs</sub><br>minus XTR | 2026 | 2026 | 2 | 4 | 857 | 857 |

### Haploid calling on X and Y in males reduces false positives compared to diploid calling but does not affect false negatives

We find that the number of false positives decreases when calling X and Y non-PARs as haploid compared to when calling them as diploid. On average, there are 17 and 3 false positives on haploid-called X and Y non-PARs, respectively - compared to 236 and 109 false positives on X and Y non-PARs, respectively, with diploid calling (**Figure 3A; Table S4**). Compared to the autosomes, the proportion of false positives to total simulated variants is substantially higher on the diploid called Y chromosome with an average proportion of false positives of 0.0378 on the Y chromosome when called as diploid compared to between 0.0007 and 0.0028 for the autosomes (**Figure 3A**). This is not observed when calling the Y chromosome as haploid - the proportion of false positives falls within autosomal range with an average value of 0.0010 (**Figure 3A**).

**Figure 3.**
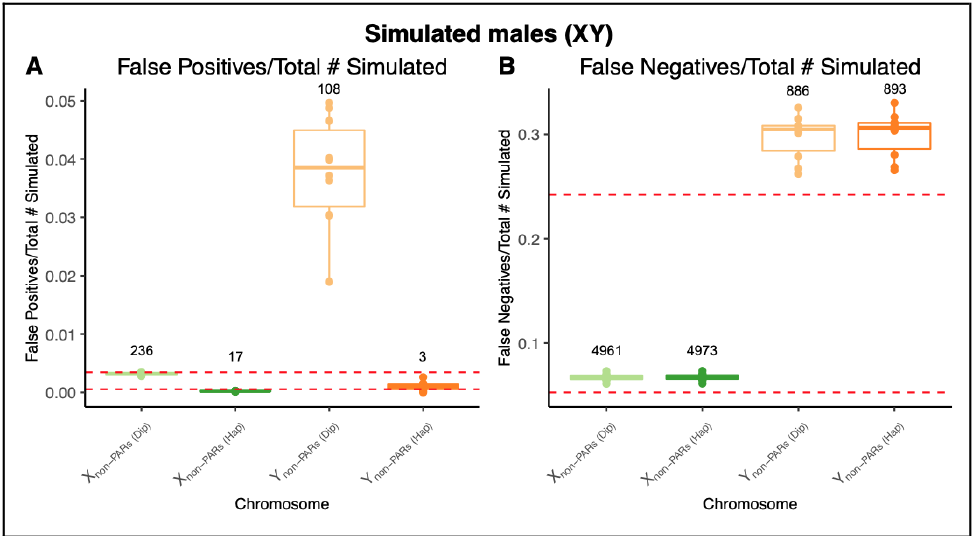
Haploid calling on X and Y in males reduces the proportion of false positives compared to diploid calling while the proportion of false negatives are similar. Box plots of A) the proportion of false positives to the total number of simulated variants and B) the proportion of false negatives to the total number of simulated variants for the sex chromosomes (X non-PARs and Y non-PARs) across 10 simulated male (XY) individuals using diploid and haploid variant calling approaches. The average number of false positives and false negatives are annotated above each box plot and highest and lowest proportions of false positives and false negatives are annotated with dashed horizontal red lines.

Unlike what was observed with false positives, the proportion of false negatives to the total number of simulated variants are roughly equal between diploid and haploid calling approaches (**Figure 3B**). There were on average 4961 and 886 false negatives on X and Y non-PARs, respectively, with diploid calling, while with haploid calling there were 4973 and 893 false negatives on X and Y non-PARs, respectively (**Figure 3B; Table S4**). The proportion of false negatives on X non-PARs fall within the autosomal range, while proportion of false negatives (to total variants called) on Y non-PARs is an order of magnitude higher than the autosomes (**Figure 3B**).

### False negatives are high across the entire Y chromosome

Overall, we find that false positives are evenly dispersed across the sex chromosomes. Proportions of false positives on the sex chromosomes are low overall and similar to autosomal ranges (**Figure S2A and S2B**). In females, the average proportion of false positives to total simulated variants is similar across the X chromosome with values of 0.0004 (4 false positives, FP), 0.0012 (126 FP), 0.0010 (1 FP), and 0.0008 (4 FP) on PAR, non-PARs, amplicons, and the XTR, respectively (**Figure 4A; Table S5**). Similarly, males have average proportions of false positives to total simulated variants of 0.0005 (3 FP), 0.0002 (16 FP), 0.0006 (<1 FP), and 0.0002 (1 FP) on PAR, non-PARs, amplicons, and the XTR, respectively (**Figure 4B; Table S5**). The proportion of false positives on Y non-PARs and amplicons are similar with average values of 0.0009 (2 FP) and 0 (0 FP) respectively. However, compared to the amplicons and rest of Y non-PARs the XTR has a slightly elevated proportion of false positives of 0.0148, though there are not many false positives in the XTR; the number of false positives range from 0 to 3 across males (**Figure 4C; Table S5**).

**Figure 4.**
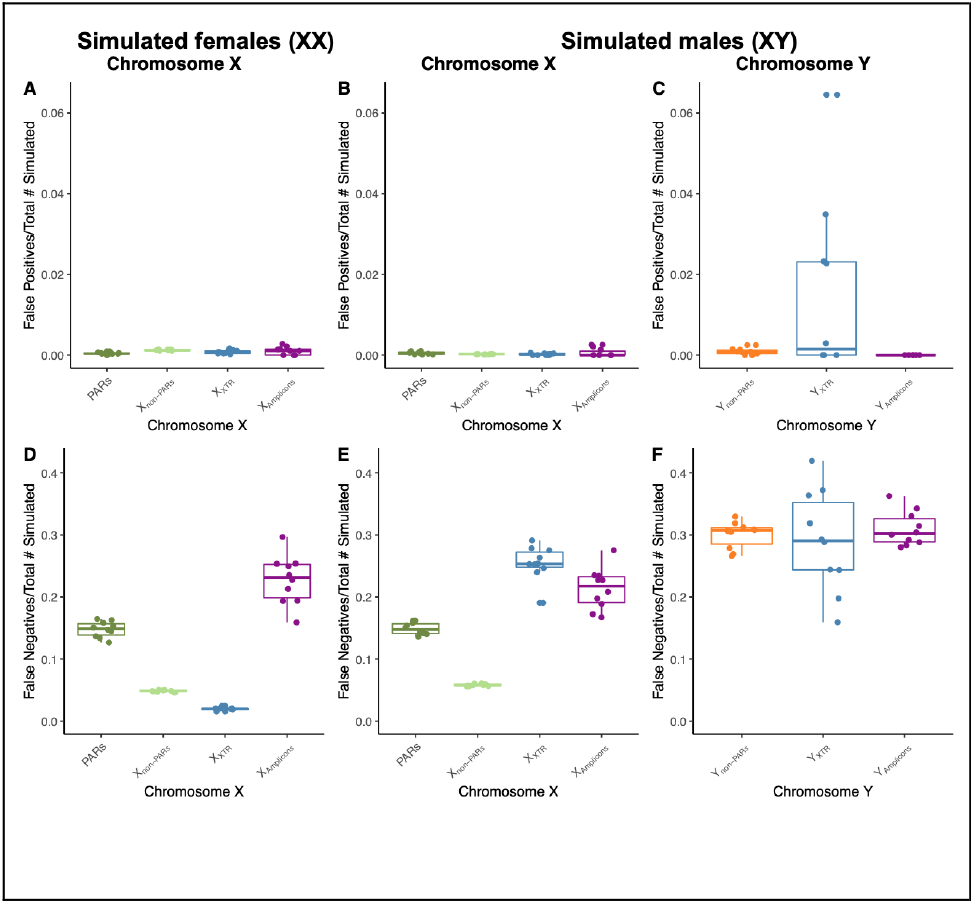
False negatives are high across all regions of the Y chromosome. Box plots of the proportion of false positives to the total number of simulated variants across A) the X chromosome in simulated females (XX), B) the X chromosome in simulated males (XY), and C) the Y chromosome in simulated males (XY), and the proportion of false negatives to the total number of simulated variants across D) the X chromosome in simulated females (XX), E) the X chromosome in simulated males (XY), and F) the Y chromosome in simulated males (XY).

Unlike false positives, we find that false negatives are elevated in regions of high sequence similarity on the X but that false negatives are evenly dispersed on the Y chromosome. Similar to what was observed for false positives, proportions of false negatives fall within autosomal ranges on the X chromosome, while false negatives have slightly higher proportions of false negatives (**Figure S2C and S2D**). On the X chromosome across simulated female individuals, the proportion of false negatives is higher on in the PARs and amplicons, with an average proportion of false negatives of 0.1480 and 0.2276, respectively, compared to non-PARs with an average proportion of false negatives of 0.0484 (**Figure 4D; Table S5**). Proportion of false negatives on the XTR is lower than X non-PARs with an average proportion of false negatives of 0.0200 (**Figure 4D; Table S5**). On the X chromosome across simulated male individuals, the proportion of false negatives is higher on in the PARs, amplicons, and XTR with average proportions of false negatives of 0.1489, 0.2133, and 0.2544, respectively, compared to 0.0582 on X non-PARs (**Figure 4E; Table S5**). In contrast, the proportions of false negatives are similar across the Y chromosome in the non-PARs, amplicons, and XTR, with an average proportion of false negatives of 0.3001, 0.3097, and 0.2899, respectively (**Figure 4F; Table S5**).

### Using diploid-based filter thresholds for some filters on haploid chromosomes is worse for variant calling

Overall, we find that using diploid-based filtering thresholds on the sex chromosomes in males for DP and AN results in a higher loss of true positives compared to haploid-based thresholds. When implementing a diploid-based DP threshold on haploid X and Y in males, there are an average of 16082 and 685 true positives on X non-PARs and Y-non-PARs, respectively (**Figure 5A; Table S6**). The number of true positives increases to an average of 58662 and 1944 true positives on X non-PARs and Y-non-PARs, respectively when implementing a haploid-based DP threshold (**Figure 5A; Table S6**). When implementing a haploid-based AN threshold of 10, there are an average of 67042 and 2333 TPs on X non-PARs and Y-non-PARs, respectively, while no true positives remain on X non-PARs and Y-non-PARs when implementing a diploid-based threshold of 20 (**Figure 5B; Table S6**).

**Figure 5.**
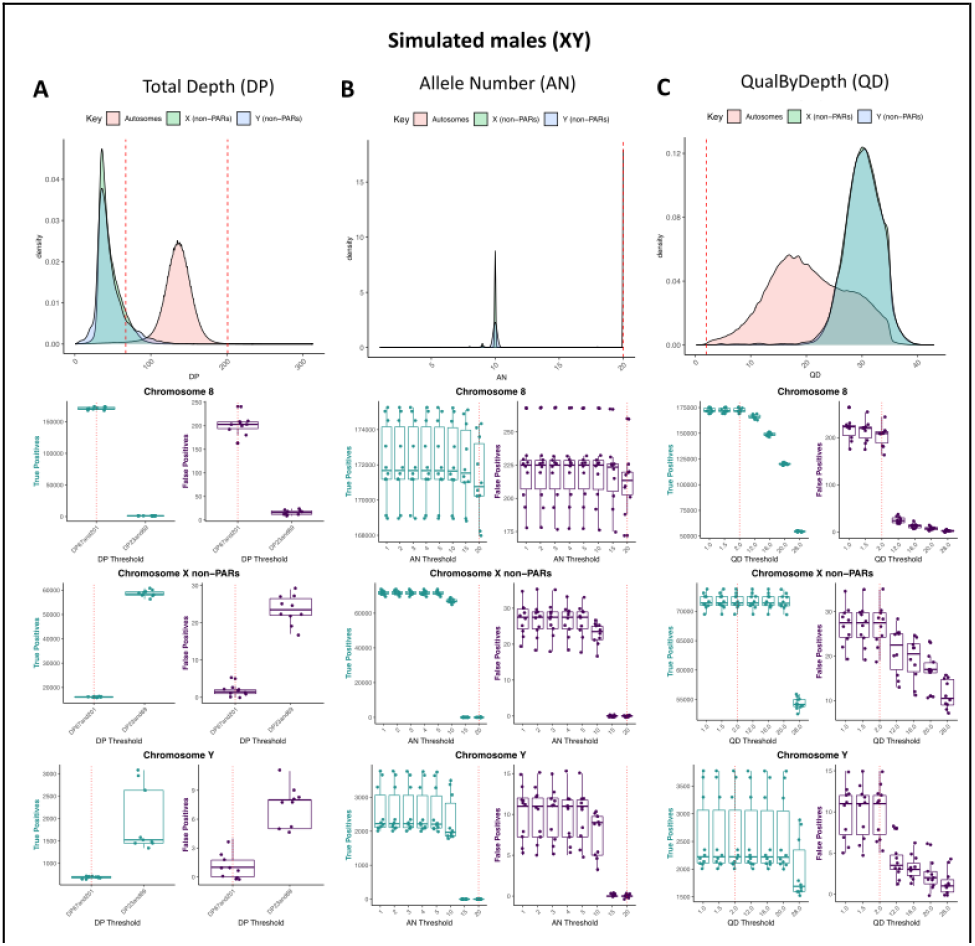
Using diploid thresholds on haploid chromosomes results in a reduction of true positives for total depth and allele number whereas increasing QualByDepth reduces the number of false positives but also reduces the number of true positives. Distributions of variants (top) and the number of true positives and false positives for different filtering thresholds for autosomes, X non-PARs, and Y non-PARs across simulated male (XY) samples for A) total depth, B) allele number, and C) QualByDepth. Chromosome 8 was used to represent the autosomes. Vertical red dashed lines represent filter thresholds based on the autosomes.

We also find that setting higher QD threshold for the sex chromosomes in males results in fewer false positives but also fewer true positives. While increasing the QD threshold from the default recommended value of 2 decreases the number of false positives from an average between 6 to 10 (for QD threshold of 12 and 20), it also results in an average loss of true positives from 4 (QD 12) to 58 (QD 20) on X non-PARs, losing more true positives much higher than the reduction in false positives. However, for the Y non-PARs, the QD threshold can be increasesd to 12 without losing any additional true positives and also decrease false positives by an average of 6 (**Figure 5C; Table S6**). In females, where the X chromosome is diploid we do not observe losses of true positives when implementing diploid DP and AN thresholds, and, similar to the autosomes, true positives start to decrease at QD threshold beyond 2 (**Figure S3; Table S6**).

### There are no large differences in the proportions of false positives and false negatives when joint genotyping across 10 or 20 samples

Overall, we find the proportion of false positives and false negatives are similar when joint genotyping across 10 and 20 samples (**Figure S4**). Across females, the proportion of false positives on the PARs is equal regardless of when joint genotyping across 10 or 20 samples (average proportion of false positives = 0.0004; **Figure S4A**). However, there is a slightly lower proportion of false positives on X non-PARs when joint genotyping 20 (average proportion of false positives = 0.0011) compared to when joint genotyping 10 samples (average proportion of false positives = 0.0012; **Figure S4A**). In males, the proportion of false positives is equal on X non-PARs (average proportion of false positives across = 0.0002). Proportion of false positives is lower on PARs in males when joint genotyping 20 (average proportion of false positives across = 0.0004) compared to when joint genotyping 10 samples (average proportion of false positives across = 0.0005; **Figure S4B**). We observed the same pattern for the Y non-PARs (average proportion of false positives across 10 samples = 0.0010; average proportion of false positives across 20 samples = 0.0008; **Figure S4B**).

The proportion of false negatives in females is slightly higher on both the PARs (average proportion of false positives across 10 samples = 0.1480; average proportion of false positives across 20 samples = 0.1502) and X non-PARs (average proportion of false positives across 10 samples = 0.0485; average proportion of false positives across 20 samples = 0.0490) when joint genotyping 20 compared to 10 samples (**Figure S4C**). Similarly, in males, the proportion of false negatives is slightly higher on the PARs (average proportion of false positives across 10 samples = 0.1489; average proportion of false positives across 20 samples = 0.1500), but slightly lower on X non-PARs (average proportion of false positives across 10 samples = 0.0673; average proportion of false positives across 20 samples = 0.0652) and Y non-PARs (average proportion of false positives across 10 samples = 0.2999; average proportion of false positives across 20 samples = 0.2881) when joint genotyping 20 compared to 10 samples (**Figure S4D**). Finally, we see no difference in the proportion of false positives or false negatives simulated across different populations (**Figure S5**).

### Clinical relevance of missed mutations

Finally, we assessed the clinical relevance of true positives that were not called when using a default alignment and variant calling strategy. In this small pilot dataset we identified 5044 variants in 45 genes on the X chromosome (**Table S7**). Within these genes, we assembled reported disease associations from the GWAS catalog (Cerezo et al. 2025), OMIM (“Home - OMIM,” n.d.), and ClinVar (Landrum et al. 2014). Based on these results, we find that these genes are implicated in multiple disease processes, including cardiomyopathy, immunodeficiency, and Alzheimer’s disease (**Figure 6; Table S7**).

**Figure 6.**
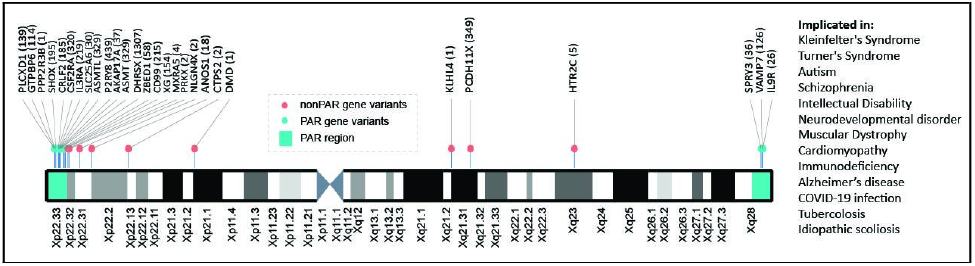
Recovery of sex chromosome variants involved in human disease. Here we highlight variants across the X chromosome that we recover using a sex chromosome aware alignment, variant calling, and filtering strategy. On the right we highlight disease processes that the genes in which these variants were identified have been implicated in.

## Discussion

Here, we undertook a simulation study to assess the effects of standard autosomal versus sex chromosome complement-informed alignment, variant calling and variant filtering strategies on variants called on the human sex chromosomes. Overall, we find that aligning samples to a reference genome informed on the sex chromosome complement of the sample, using biological relevant ploidy for calling variants, and setting filtering thresholds informed by ploidy of the chromosome improves variant calling on the sex chromosomes. We found more true positives on the X chromosome when aligning samples to a sex chromosome complement reference genome compared to a default reference genome. This suggests that implementing sex chromosome complement-informed alignment improves variant calling compared to a standard default alignment. Previously, it has been reported that more variants are called on the X chromosome in female samples when aligning to a sex chromosome complement reference (Webster et al. 2019). However, it was not confirmed whether these additional variants were in fact true positives and whether this was observed in male samples. We expand on this work by using simulated sequence data in both female and male samples, finding more true positives in PARs in males and females and the XTR in females when aligning to a sex chromosome complement reference compared to a default reference genome. We found no differences in true positives on the Y non-PARs. This is not surprising, as the Y non-PAR sequence was not altered in the alignment to male samples.

Surprisingly, we observed a ten-fold higher rate of false positives in the XTR when masking Y-linked XTR during alignment of male samples, suggesting that masking one copy of the XTR reduces the accuracy of variant calling in the XTR in males. The XTR between X and Y is very similar, with 98.78% homology (Veerappa, Padakannaya, and Ramachandra 2013), so by forcing the reads to one chromosome - in this case the X chromosome - we expected variant calling to improve in XTR. However, this was not what was observed and we therefore do not recommend masking the XTR during the alignment of male samples. These results highlight the need for a Y-specific alignment approach in the future.

Currently, the human reference genomes available do not provide sex chromosome complement versions, nor directions for alternatively mapping approaches for the sex chromosomes. For example, Ensembl’s human reference does mask the PARs on the Y chromosome, which would be appropriate for aligning males. However, when aligning samples without a Y chromosome, the entire Y chromosome should be hard masked. The GENCODE human reference genome contains full sequences of the X and Y chromosomes (Schneider et al. 2017). Additionally, the Telomere-to-Telomere genome (T2T-CHM13) has recently been assembled (Nurk et al. 2022). T2T-CHM13 does not contain the Y chromosome but the newest deposited version contains a Y chromosome from HG002 (GenBank assembly accession: GCA_009914755.4). We anticipate wide use of the telomere-to-telomere resource in the near future as it is an improvement from the GRCh38 reference genome (Nurk et al. 2022). Tools like XYalign or software like bedtools maskfasta can facilitate generation of sex chromosome complement versions of any of these reference genomes (Webster et al. 2019; Quinlan and Hall 2010).

In terms of variant calling strategy, we found that haploid calling on the sex chromosomes in males reduces false positives compared to diploid calling but does not affect false negatives. Here, we stress the importance of biologically accurate ploidy when calling variants to reduce the chance of identifying false associations with disease, or falsely inferring demography from Y chromosome variants. When using GATK, ploidy is specified when generating GVCFs using HaplotypeCaller (Poplin et al., n.d.). During the joint genotyping step, the GenotypeGVCFs tool is able to handle mixed ploidy, so it is possible to joint call, for example, the X chromosome across both male and female samples (where the X chromosome is haploid in males and diploid in females).

We also found that using diploid-based filtering thresholds reduces variant calling accuracy on the sex chromosomes in males. Specifically, we found that implementing diploid-based thresholds for DP and AN results in a higher loss of true positives compared to haploid-based thresholds. The sex chromosomes in males are haploid, so both DP and AN should be about half of diploid chromosomes. If processing the sex chromosomes like autosomes, we expect that DP and AN thresholds with diploid settings would reduce the number of true positives on X and Y chromosomes in males compared to thresholds optimized for haploid chromosomes, as these diploid-based thresholds are likely too stringent for haploid chromosomes. Additionally, since QD is inversely based on DP, we expected to retain true positives for higher QD thresholds for X and Y in males compared to the autosomes. However, though setting higher QD threshold for the sex chromosomes in males results in fewer false positives, we also observed a reduction in true positives. Previous studies have highlighted the importance of filtering variants to retain an accurate call set (e.g. (Jia et al. 2012; Reumers et al. 2011; DePristo et al. 2011)). Here, we additionally highlight the importance of considering ploidy when optimizing filtering thresholds on the sex chromosomes. We further find that proportions of false positives and negatives are similar on the X and Y in females and males when joint genotyping additional samples or processing simulated reads from individuals of different ancestries (African, Asian, and European).

Finally, we observe that genes with newly-recovered variants have been implicated in several human disease processes. However, a concern remains, that there may be a vast underrepresentation of sex chromosome-linked disease variants due to a historical exclusion of the X in genome-wide association studies (Sun et al. 2023), as well as not even identifying the variants that would be needed to be included in the study (Pinto et al. 2023b).

## Conclusion

Here, we assessed the effects of implementing standard autosomal versus sex chromosome complement-informed alignment, variant calling and variant filtering strategies on variants called on the human sex chromosomes and provide recommendations for mapping, calling and filtering variants on the sex chromosomes. We find that aligning to a reference genome informed on the sex chromosome complement of samples improves variant calling on the sex chromosome compared to aligning to a default reference, and variant calling is improved in males when calling the sex chromosomes haploid rather than diploid and when using haploid-based thresholds for filtering variants on the sex chromosomes. We recommend aligning samples to versions of the human reference genome informed on the sex chromosome complement of the sample and to use biologically accurate ploidy parameters when calling variants and setting filtering thresholds. This approach will help improve future genetic studies that include the sex chromosomes.

## Data and code availability

All original code used in this manuscript can be found on GitHub: https://github.com/SexChrLab/VariantCalling.

## Acknowledgements

The authors acknowledge Research Computing at Arizona State University for providing high performance computing resources that have contributed to the research results reported within this paper.

## Funding

This work was supported, in part, by the Intramural Research Program of the National Human Genome Research Institute, National Institutes of Health. This publication was supported by the National Institute of General Medical Sciences of the National Institutes of Health under Award Number R35GM124827 to MAW.

## Notes

https://github.com/SexChrLab/VariantCalling

### Competing Interest Statement

The authors have declared no competing interest.

