## Supplementary Material for "Best practices for improving alignment and variant calling on human sex chromosomes"

### Supplementary Materials

**Supplementary Tables** **2**

Table S1. Aligning reads to sex chromosome complement reference genome improves variant calling on the X chromosome compared to aligning to a default reference genome. 2

Table S2. Per sample performance metrics on X and Y. 4

Table S3. Genomic features on X and Y. 8

Table S2. Variants were introduced into simulated sequences from male individuals from 1000 Genomes. 9

Table S3. Effects of masking the X-transposed region (XTR). 13

Table S4. Effects of haploid calling. 15

Table S5. Counts of variants across regions of the X and Y. 16

Table S6. Using diploid-based filter thresholds for some filters on haploid chromosomes results in less accurate variant calling. 20

**Supplementary Figures** **22**

Supplementary Figure 1. Variant calling accuracy declines substantially in XTR when masking XTR on the Y chromosome for alignment. 22

Figure S2. The proportion of false positives and false negatives on chromosome X are similar to autosomal ranges while chromosome Y has an elevated proportion of false negatives. 23

Figure S3. Distribution of DP, AN, and QD are similar between autosomes and the X chromosome in females 24

### Supplementary Tables

#### Table S1. Aligning reads to sex chromosome complement reference genome improves variant calling on the X chromosome compared to aligning to a default reference genome.

Average number of true positives, false positives, and false negatives on the X chromosome across 10 simulated female (XX) samples and 10 simulated male (XY) samples and across the Y chromosome in 10 simulated male (XY) samples using a default and sex chromosome complement alignment approaches. Chromosome X was split into PARs, XTR, and non-PARs without XTR and chromosome Y was split into XTR, and non-PARs without XTR.

| Chr | Region | Joint Genotyping with | Average # TP | | Average # FP | | Average # FN | |
|---|---|---|---|---|---|---|---|---|
| | | | Default | SCC | Default | SCC | Default | SCC |
| X | PARs | 10 Females (XX) | 0 | 4909 | 0 | 2 | 5763 | 853 |
| | | 10 Males (XY) | 0 | 4980 | 0 | 3 | 5852 | 872 |
| | | 10 Females (XX) + 10 Males (XY) | 0 | 4939 (4910 avg F, 4969 avg M) | 0 | 2 (2 avg F, 2 avg M) | 5807 (5763 avg F, 5852 avg M) | 867 (852 avg F, 882 avg M) |
| | | 20 Females (XX) | 0 | 4947 | 0 | 2 | 5823 | 875 |
| | | 20 Males (XY) | 1 | 5105 | 0 | 2 | 6004 | 900 |
| | | 20 Females (XX) + 20 Males (XY) | 1 (0 avg F, 1 avg M) | 5013 (4934 avg F, 5091 avg M) | 0 | 2 (2 avg F, 2 avg M) | 5914 (5823 avg F, 6004 avg M) | 901 (888 avg F, 913 avg M) |
| | non PARs (minus XTR) | 10 Females (XX) | 97994 | 98007 | 127 | 127 | 5148 | 5135 |
| | | 10 Males (XY) | 66780 | 66780 | 17 | 16 | 4217 | 4217 |
| | | 10 Females (XX) + 10 Males (XY) | 82840 (98063 avg F, 67618 avg M) | 82851 (98007 avg F, 67626 avg M) | 63 (115 avg F, 11 avg M) | 63 (115 avg F, 11 avg M) | 4228 (5077 avg F, 3379 avg M) | 4217 (5062 avg F, 3371 avg M) |
| | | 20 Females (XX) | 97778 | 97789 | 114 | 114 | 5194 | 5183 |
| | | 20 Males (XY) | 66745 | 66746 | 17 | 17 | 4053 | 4051 |
| | | 20 Females (XX) + 20 Males (XY) | 82615 (97810 avg F, 67420 avg M) | 82627 (97826 avg F, 67428avg M) | 55 (99 avg F, 11 avg M) | 55 (99 avg F, 11 avg M) | 4267 (5159 avg F, 3376 avg M) | 4256 (5143 avg F, 3368 avg M) |
| | XTR | 10 Females (XX) | 3767 | 4515 | 3 | 4 | 841 | 93 |
| | | 10 Males (XY) | 2171 | 2171 | 1 | 1 | 756 | 756 |
| | | 10 Females (XX) + 10 Males (XY) | 3043 (3744 F, | 3586 (4513 F, | 1 (2 F, 0 M) | 2 (3 F, 0 M) | 725 (864 F, 586 M) | 181 (94 F, 268 |

| | | | | | | | | |
|---|---|---|---|---|---|---|---|---|
| | | | 2342 M) | 2659 M) | | | | M) |
| | | 20 Females (XX) | 3834 | 4563 | 3 | 3 | 818 | 89 |
| | | 20 Males (XY) | 2202 | 2202 | 0 | 0 | 761 | 761 |
| | | 20 Females (XX) + 20 Males (XY) | 3094 (3805 avg F, 2383 avg M) | 3636 (4562 avg F, 2711 avg M) | 1 (2 avg F, 0 avg M) | 1 (2 avg F, 0 avg M) | 713 (847 avg F, 560 avg M) | 171 (90 avg F, 252 avg M) |
| Y | non PARs (minus XTR) | 10 Males (XY) | 2026 | 2026 | 2 | 2 | 857 | 857 |
| | | 20 Males (XY) | 2262 | 2262 | 2 | 2 | 902 | 902 |
| | XTR | 10 Males (XY) | 103 | 103 | 1 | 1 | 36 | 36 |
| | | 20 Males (XY) | 138 | 138 | 1 | 1 | 51 | 51 |

### Table S2. Per sample performance metrics on X and Y.

This is for 20 males and 20 females. 20 males were joint genotyped together and 20 females were joint genotyped together separately. Male X and Y non PARs called haploid. SCC aligned data.

| Region | Sample size | Sample | # True Positives | # False Positives | # False Negatives | # Simulated |
|---|---|---|---|---|---|---|
| PARs | 20 Males (XY) | NA06984 | 4639 | 1 | 915 | 5554 |
| | | NA06986 | 4921 | 0 | 815 | 5736 |
| | | NA06994 | 5123 | 1 | 823 | 5946 |
| | | NA07048 | 5205 | 6 | 947 | 6152 |
| | | NA07051 | 5016 | 4 | 986 | 6002 |
| | | NA07347 | 5031 | 1 | 941 | 5972 |
| | | NA07357 | 4722 | 1 | 795 | 5517 |
| | | NA10851 | 5042 | 6 | 850 | 5892 |
| | | NA11829 | 4983 | 2 | 954 | 5937 |
| | | NA11831 | 4959 | 1 | 854 | 5813 |
| | | NA11843 | 5297 | 1 | 850 | 6147 |
| | | NA11881 | 5214 | 3 | 812 | 6026 |
| | | NA11893 | 5293 | 6 | 982 | 6275 |
| | | NA11919 | 5253 | 0 | 929 | 6182 |
| | | NA11930 | 5115 | 0 | 831 | 5946 |
| | | NA11932 | 5138 | 2 | 882 | 6020 |
| | | NA11992 | 5468 | 1 | 953 | 6421 |
| | | NA11994 | 5594 | 3 | 930 | 6524 |
| | | NA12003 | 5130 | 2 | 985 | 6115 |
| | | NA12005 | 4950 | 2 | 975 | 5925 |
| | 20 Females (XX) | NA06985 | 5218 | 3 | 942 | 6161 |
| | | NA06989 | 4760 | 1 | 938 | 5699 |
| | | NA07000 | 5225 | 2 | 835 | 6061 |
| | | NA07037 | 5064 | 2 | 738 | 5803 |
| | | NA07056 | 5036 | 2 | 858 | 5895 |
| | | NA10847 | 4975 | 1 | 891 | 5867 |
| | | NA11830 | 4604 | 2 | 713 | 5318 |
| | | NA11832 | 4638 | 0 | 895 | 5533 |
| | | NA11840 | 5013 | 1 | 952 | 5966 |
| | | NA11892 | 4552 | 4 | 773 | 5326 |
| | | NA11894 | 5172 | 6 | 798 | 5971 |
| | | NA11918 | 5105 | 1 | 1072 | 6178 |
| | | NA11920 | 5015 | 2 | 921 | 5936 |
| | | NA11931 | 4693 | 2 | 911 | 5605 |
| | | NA11933 | 5003 | 1 | 715 | 5719 |
| | | NA11995 | 5170 | 3 | 934 | 6105 |
| | | NA12004 | 5114 | 2 | 859 | 5973 |
| | | NA12006 | 4666 | 1 | 902 | 5569 |
| | | NA12044 | 4890 | 1 | 897 | 5789 |

| | | | | | | |
|---|---|---|---|---|---|---|
| | | NA12046 | 5028 | 5 | 965 | 5994 |
| X non PARs (minus XTR) | 20 Males (XY) | NA06984 | 67367 | 22 | 4087 | 71458 |
| | | NA06986 | 68227 | 20 | 4016 | 72246 |
| | | NA06994 | 66423 | 20 | 4237 | 70662 |
| | | NA07048 | 64626 | 14 | 3788 | 68419 |
| | | NA07051 | 66676 | 13 | 3963 | 70643 |
| | | NA07347 | 66338 | 20 | 4238 | 70579 |
| | | NA07357 | 68635 | 13 | 4350 | 72989 |
| | | NA10851 | 67037 | 10 | 3956 | 70997 |
| | | NA11829 | 65732 | 14 | 3972 | 69708 |
| | | NA11831 | 68217 | 16 | 4070 | 72291 |
| | | NA11843 | 68996 | 19 | 4294 | 73294 |
| | | NA11881 | 66046 | 13 | 4088 | 70138 |
| | | NA11893 | 67934 | 19 | 3993 | 71931 |
| | | NA11919 | 68243 | 16 | 4004 | 72251 |
| | | NA11930 | 64872 | 26 | 3811 | 68687 |
| | | NA11932 | 68634 | 14 | 3943 | 72579 |
| | | NA11992 | 64121 | 17 | 4110 | 68235 |
| | | NA11994 | 65956 | 17 | 4073 | 70033 |
| | | NA12003 | 64990 | 16 | 4113 | 69108 |
| | | NA12005 | 65858 | 16 | 3920 | 69781 |
| | Females (XX) | NA06985 | 96616 | 116 | 4854 | 101473 |
| | | NA06989 | 98587 | 116 | 5144 | 103736 |
| | | NA07000 | 100040 | 104 | 5461 | 105505 |
| | | NA07037 | 98692 | 118 | 5354 | 104053 |
| | | NA07056 | 99161 | 118 | 5108 | 104274 |
| | | NA10847 | 97279 | 109 | 5077 | 102362 |
| | | NA11830 | 95996 | 124 | 4894 | 100895 |
| | | NA11832 | 98222 | 99 | 5392 | 103618 |
| | | NA11840 | 99263 | 125 | 4976 | 104244 |
| | | NA11892 | 96104 | 116 | 5166 | 101272 |
| | | NA11894 | 100663 | 100 | 5244 | 105913 |
| | | NA11918 | 96290 | 129 | 5152 | 101447 |
| | | NA11920 | 97972 | 123 | 5481 | 103458 |
| | | NA11931 | 96058 | 107 | 5484 | 101548 |
| | | NA11933 | 95590 | 103 | 5225 | 100820 |
| | | NA11995 | 98925 | 89 | 5210 | 104141 |
| | | NA12004 | 100001 | 117 | 5490 | 105495 |
| | | NA12006 | 96246 | 115 | 4760 | 101014 |
| | | NA12044 | 98269 | 123 | 5165 | 103437 |
| | | NA12046 | 95804 | 127 | 5028 | 100837 |
| X XTR | 20 Males (XY) | NA06984 | 2490 | 1 | 874 | 3364 |
| | | NA06986 | 2243 | 0 | 755 | 2998 |
| | | NA06994 | 1795 | 1 | 570 | 2365 |
| | | NA07048 | 2415 | 0 | 791 | 3206 |
| | | NA07051 | 2029 | 0 | 673 | 2702 |
| | | NA07347 | 2498 | 0 | 1026 | 3525 |

| | | | | | | |
|---|---|---|---|---|---|---|
| | | NA07357 | 2441 | 1 | 900 | 3341 |
| | | NA10851 | 1653 | 0 | 371 | 2024 |
| | | NA11829 | 2483 | 2 | 937 | 3420 |
| | | NA11831 | 1760 | 1 | 568 | 2328 |
| | | NA11843 | 2370 | 0 | 911 | 3281 |
| | | NA11881 | 2217 | 0 | 736 | 2953 |
| | | NA11893 | 2192 | 0 | 860 | 3052 |
| | | NA11919 | 2272 | 0 | 671 | 2943 |
| | | NA11930 | 2503 | 1 | 1010 | 3514 |
| | | NA11932 | 2297 | 0 | 752 | 3049 |
| | | NA11992 | 2240 | 0 | 727 | 2968 |
| | | NA11994 | 1793 | 0 | 570 | 2363 |
| | | NA12003 | 2225 | 0 | 790 | 3016 |
| | | NA12005 | 2127 | 1 | 725 | 2852 |
| | | NA06985 | 4419 | 2 | 61 | 4480 |
| | | NA06989 | 4573 | 4 | 93 | 4666 |
| | | NA07000 | 4752 | 6 | 100 | 4853 |
| | | NA07037 | 4859 | 3 | 97 | 4956 |
| | | NA07056 | 4535 | 2 | 93 | 4629 |
| | | NA10847 | 4789 | 2 | 112 | 4901 |
| | | NA11830 | 3982 | 4 | 82 | 4064 |
| | | NA11832 | 4512 | 1 | 84 | 4596 |
| | | NA11840 | 3775 | 2 | 73 | 3848 |
| | Females (XX) | NA11892 | 4992 | 3 | 91 | 5083 |
| | | NA11894 | 4707 | 4 | 97 | 4804 |
| | | NA11918 | 5157 | 7 | 115 | 5273 |
| | | NA11920 | 4549 | 3 | 84 | 4633 |
| | | NA11931 | 5371 | 5 | 93 | 5465 |
| | | NA11933 | 4307 | 3 | 77 | 4384 |
| | | NA11995 | 4375 | 2 | 76 | 4451 |
| | | NA12004 | 4779 | 2 | 103 | 4882 |
| | | NA12006 | 4364 | 1 | 70 | 4435 |
| | | NA12044 | 4230 | 2 | 82 | 4312 |
| | | NA12046 | 4236 | 5 | 90 | 4326 |
| | | NA06984 | 1761 | 1 | 783 | 2544 |
| | | NA06986 | 1953 | 4 | 833 | 2787 |
| | | NA06994 | 2745 | 2 | 1053 | 3799 |
| | | NA07048 | 1812 | 3 | 795 | 2608 |
| | | NA07051 | 3144 | 4 | 1132 | 4277 |
| | | NA07347 | 1691 | 3 | 806 | 2498 |
| Y non PARs | 20 Males (XY) | NA07357 | 1852 | 7 | 817 | 2669 |
| | | NA10851 | 1712 | 2 | 763 | 2475 |
| | | NA11829 | 3056 | 2 | 1099 | 4156 |
| | | NA11831 | 1656 | 0 | 751 | 2407 |
| | | NA11843 | 2396 | 2 | 905 | 3302 |
| | | NA11881 | 2961 | 2 | 1101 | 4062 |
| | | NA11893 | 2721 | 1 | 1075 | 3797 |

| | | | | | | |
|---|---|---|---|---|---|---|
| | | NA11919 | 2999 | 5 | 1128 | 4128 |
| | | NA11930 | 2898 | 2 | 1034 | 3933 |
| | | NA11932 | 2970 | 3 | 1112 | 4082 |
| | | NA11992 | 2792 | 3 | 1105 | 3898 |
| | | NA11994 | 1852 | 1 | 801 | 2653 |
| | | NA12003 | 3136 | 3 | 1102 | 4239 |
| | | NA12005 | 1894 | 2 | 863 | 2758 |

### Table S3. Genomic features on X and Y.

All analyses in this manuscript were performed in GRCh38. We lifted over XTR and amplicons using the UCSC liftover tool. PARs were obtained from Ensembl. X chromosome ampliconic region 3 failed liftover and ampliconic region 5 split into 4 regions.

| Chromosome | Region | Coordinate (GRCh37/hg19) | Coordinate (GRCh38) |
|---|---|---|---|
| X | PAR1 | 60001 - 2699520 | 10001 - 2781479 |
| | PAR2 | 154931044 - 155260560 | 155701383 - 156030895 |
| | XTR | 88193855 - 93193855 | 89140830 - 93428068 |
| | Ampliconic region 1 | 48202745 - 48292983 | 48343310 - 48434604 |
| | Ampliconic region 2 | 48976199 - 49062381 | 49157254 - 49162986 |
| | Ampliconic region 3 | 51395467 - 51492862 | N/A |
| | Ampliconic region 4 | 51775560 - 51966529 | 52032464 - 52223402 |
| | Ampliconic region 5 | 52518132 - 53027386 | 52511000 - 52511492 |
| | | | 52489200 - 52501394 |
| | | | 52520509 - 52520537 |
| | | | 139870917 - 139871081 |
| | Ampliconic region 6 | 55464117 - 55574172 | 55437684 - 55547739 |
| | Ampliconic region 7 | 62335733 - 62495350 | 63185573 - 63193371 |
| | Ampliconic region 8 | 70894117 - 71055682 | 71674267 - 71835832 |
| | Ampliconic region 9 | 71941159 - 72325075 | 72721314 - 73105236 |
| | Ampliconic region 10 | 100818723 - 100903977 | 101563740 - 101648990 |
| | Ampliconic region 11 | 101435778 - 101774391 | 102180805 - 102519463 |
| | Ampliconic region 12 | 103195105 - 103362341 | 103940531 - 104117650 |
| Y | PAR1 | 10001 - 2649520 | 10001 - 2781479 |
| | PAR2 | 59034050 - 59363566 | 56987321 - 57217415 |
| | XTR1 | 2918085 - 6103152 | 3050044 - 6235111 |
| | XTR2 | 6400947 - 6616754 | 6532906 - 6748713 |
| | Ampliconic region 1 | 6103152 - 6400947 | 6235111 - 6532906 |
| | Ampliconic region 2 | 7442522 - 10135224 | 7574481 - 10266944 |
| | Ampliconic region 3 | 16096353 - 16170613 | 13984473 - 14058733 |
| | Ampliconic region 4 | 17986973 - 18017094 | 15875093 - 15905214 |
| | Ampliconic region 5 | 18271675 - 18537845 | 16159795 - 16425965 |
| | Ampliconic region 6 | 19568146 - 21032220 | 17456266 - 18870334 |
| | Ampliconic region 7 | 23467839 - 28561582 | 21305953 - 26415435 |

### Table S2. Variants were introduced into simulated sequences from male individuals from 1000 Genomes.

Variants were introduced into simulated sequence reads using variant call data from 1000 Genomes males and females from individuals of European, African, and Asian descent.

| Sample ID | Population | Super population | Sex |
|---|---|---|---|
| NA06984 | CEU | EUR | Male |
| NA06986 | CEU | EUR | Male |
| NA06994 | CEU | EUR | Male |
| NA07048 | CEU | EUR | Male |
| NA07051 | CEU | EUR | Male |
| NA07347 | CEU | EUR | Male |
| NA07357 | CEU | EUR | Male |
| NA10851 | CEU | EUR | Male |
| NA11829 | CEU | EUR | Male |
| NA11831 | CEU | EUR | Male |
| NA11843 | CEU | EUR | Male |
| NA11881 | CEU | EUR | Male |
| NA11893 | CEU | EUR | Male |
| NA11919 | CEU | EUR | Male |
| NA11930 | CEU | EUR | Male |
| NA11932 | CEU | EUR | Male |
| NA11992 | CEU | EUR | Male |
| NA11994 | CEU | EUR | Male |
| NA12003 | CEU | EUR | Male |
| NA12005 | CEU | EUR | Male |
| NA06985 | CEU | EUR | Female |
| NA06989 | CEU | EUR | Female |
| NA07000 | CEU | EUR | Female |
| NA07037 | CEU | EUR | Female |
| NA07056 | CEU | EUR | Female |
| NA10847 | CEU | EUR | Female |
| NA11830 | CEU | EUR | Female |
| NA11832 | CEU | EUR | Female |
| NA11840 | CEU | EUR | Female |
| NA11892 | CEU | EUR | Female |
| NA11894 | CEU | EUR | Female |
| NA11918 | CEU | EUR | Female |

| | | | |
|---|---|---|---|
| NA11920 | CEU | EUR | Female |
| NA11931 | CEU | EUR | Female |
| NA11933 | CEU | EUR | Female |
| NA11995 | CEU | EUR | Female |
| NA12004 | CEU | EUR | Female |
| NA12006 | CEU | EUR | Female |
| NA12044 | CEU | EUR | Female |
| NA12046 | CEU | EUR | Female |
| NA18486 | YRI | AFR | Male |
| NA18498 | YRI | AFR | Male |
| NA18501 | YRI | AFR | Male |
| NA18504 | YRI | AFR | Male |
| NA18507 | YRI | AFR | Male |
| NA18510 | YRI | AFR | Male |
| NA18516 | YRI | AFR | Male |
| NA18519 | YRI | AFR | Male |
| NA18522 | YRI | AFR | Male |
| NA18853 | YRI | AFR | Male |
| NA18856 | YRI | AFR | Male |
| NA18865 | YRI | AFR | Male |
| NA18868 | YRI | AFR | Male |
| NA18871 | YRI | AFR | Male |
| NA18874 | YRI | AFR | Male |
| NA18877 | YRI | AFR | Male |
| NA18879 | YRI | AFR | Male |
| NA18908 | YRI | AFR | Male |
| NA18910 | YRI | AFR | Male |
| NA18915 | YRI | AFR | Male |
| NA18488 | YRI | AFR | Female |
| NA18489 | YRI | AFR | Female |
| NA18499 | YRI | AFR | Female |
| NA18502 | YRI | AFR | Female |
| NA18505 | YRI | AFR | Female |
| NA18508 | YRI | AFR | Female |
| NA18511 | YRI | AFR | Female |
| NA18517 | YRI | AFR | Female |
| NA18520 | YRI | AFR | Female |
| NA18523 | YRI | AFR | Female |

| | | | |
|---|---|---|---|
| NA18858 | YRI | AFR | Female |
| NA18861 | YRI | AFR | Female |
| NA18864 | YRI | AFR | Female |
| NA18867 | YRI | AFR | Female |
| NA18870 | YRI | AFR | Female |
| NA18873 | YRI | AFR | Female |
| NA18876 | YRI | AFR | Female |
| NA18878 | YRI | AFR | Female |
| NA18881 | YRI | AFR | Female |
| NA18907 | YRI | AFR | Female |
| NA18530 | CHB | EAS | Male |
| NA18534 | CHB | EAS | Male |
| NA18536 | CHB | EAS | Male |
| NA18543 | CHB | EAS | Male |
| NA18544 | CHB | EAS | Male |
| NA18546 | CHB | EAS | Male |
| NA18548 | CHB | EAS | Male |
| NA18549 | CHB | EAS | Male |
| NA18557 | CHB | EAS | Male |
| NA18558 | CHB | EAS | Male |
| NA18559 | CHB | EAS | Male |
| NA18561 | CHB | EAS | Male |
| NA18562 | CHB | EAS | Male |
| NA18563 | CHB | EAS | Male |
| NA18572 | CHB | EAS | Male |
| NA18603 | CHB | EAS | Male |
| NA18605 | CHB | EAS | Male |
| NA18606 | CHB | EAS | Male |
| NA18608 | CHB | EAS | Male |
| NA18609 | CHB | EAS | Male |
| NA18525 | CHB | EAS | Female |
| NA18526 | CHB | EAS | Female |
| NA18528 | CHB | EAS | Female |
| NA18531 | CHB | EAS | Female |
| NA18532 | CHB | EAS | Female |
| NA18533 | CHB | EAS | Female |
| NA18535 | CHB | EAS | Female |
| NA18537 | CHB | EAS | Female |

| | | | |
|---|---|---|---|
| NA18538 | CHB | EAS | Female |
| NA18539 | CHB | EAS | Female |
| NA18541 | CHB | EAS | Female |
| NA18542 | CHB | EAS | Female |
| NA18545 | CHB | EAS | Female |
| NA18547 | CHB | EAS | Female |
| NA18550 | CHB | EAS | Female |
| NA18552 | CHB | EAS | Female |
| NA18553 | CHB | EAS | Female |
| NA18555 | CHB | EAS | Female |
| NA18560 | CHB | EAS | Female |
| NA18564 | CHB | EAS | Female |

### Table S3. Effects of masking the X-transposed region (XTR).

Masking XTR in the reference genome increases the number of false positives in XTR but does not impact variant calling on the rest of the X and Y chromosomes. Number of true positives, false positives, and false negatives in XTR, X non-PARs (excluding XTR), and Y non-PARs (excluding XTR) in males when aligning to a reference genome with just the PARs masked on the Y chromosome and PARs and XTR masked on the Y chromosome.

| **Region** | **Sample** | **# TPs** | | **# FPs** | | **# FNs** | |
|---|---|---|---|---|---|---|---|
| | | **Y PARs masked** | **Y PARs + XTR masked** | **Y PARs masked** | **Y PARs + XTR masked** | **Y PARs masked** | **Y PARs + XTR masked** |
| XTR | NA11831 | 1754 | 2228 | 1 | 33580 | 574 | 75 |
| | NA11829 | 2468 | 3300 | 2 | 33180 | 952 | 88 |
| | NA10851 | 1638 | 1934 | 0 | 33808 | 386 | 76 |
| | NA07357 | 2422 | 3222 | 1 | 33113 | 919 | 85 |
| | NA07347 | 2499 | 3366 | 0 | 33042 | 1026 | 128 |
| | NA07051 | 2016 | 2554 | 0 | 33466 | 686 | 130 |
| | NA07048 | 2395 | 3061 | 0 | 33302 | 811 | 119 |
| | NA06994 | 1798 | 2271 | 1 | 33479 | 567 | 63 |
| | NA06986 | 2241 | 2853 | 0 | 33335 | 757 | 113 |
| | NA06984 | 2478 | 3220 | 1 | 33144 | 886 | 107 |
| $X_{\text{non-PARs minus XTR}}$ | NA11831 | 68072 | 68070 | 17 | 17 | 4216 | 4218 |
| | NA11829 | 65576 | 65576 | 13 | 13 | 4130 | 4130 |
| | NA10851 | 66915 | 66917 | 10 | 10 | 4080 | 4078 |
| | NA07357 | 68508 | 68508 | 17 | 17 | 4479 | 4479 |
| | NA07347 | 66168 | 66170 | 20 | 20 | 4410 | 4408 |
| | NA07051 | 66517 | 66517 | 14 | 14 | 4124 | 4124 |
| | NA07048 | 64505 | 64500 | 13 | 14 | 3910 | 3915 |
| | NA06994 | 66257 | 66254 | 19 | 19 | 4403 | 4406 |
| | NA06986 | 68067 | 68067 | 22 | 22 | 4177 | 4177 |
| | NA06984 | 67216 | 67214 | 19 | 19 | 4240 | 4242 |
| $Y_{\text{non-PARs minus XTR}}$ | NA11831 | 1616 | 1616 | 0 | 1 | 750 | 750 |
| | NA11829 | 2793 | 2793 | 0 | 1 | 1021 | 1021 |
| | NA10851 | 1680 | 1680 | 1 | 2 | 752 | 752 |
| | NA07357 | 1810 | 1810 | 6 | 7 | 815 | 815 |
| | NA07347 | 1625 | 1625 | 3 | 4 | 803 | 803 |
| | NA07051 | 2873 | 2873 | 5 | 6 | 1067 | 1067 |
| | NA07048 | 1717 | 1717 | 2 | 4 | 777 | 777 |
| | NA06994 | 2533 | 2533 | 2 | 5 | 984 | 984 |

| | | | | | | | |
|---|---|---|---|---|---|---|---|
| | NA06986 | 1871 | 1871 | 1 | 2 | 829 | 829 |
| | NA06984 | 1743 | 1743 | 2 | 3 | 770 | 770 |

### Table S4. Effects of haploid calling.

Haploid calling on X and Y in males reduces false positives compared to diploid calling but does not affect false negatives. Performance metrics in X and Y non-PARs for 10 simulated male samples when variant calling in diploid and haploid modes.

| Region | Sample | # TPs | | # FPs | | # FNs | |
|---|---|---|---|---|---|---|---|
| | | Diploid | Haploid | Diploid | Haploid | Diploid | Haploid |
| $X_{non\text{-}PARs}$ | NA11831 | 69836 | 69826 | 252 | 18 | 4780 | 4790 |
| | NA11829 | 68060 | 68044 | 246 | 15 | 5066 | 5082 |
| | NA10851 | 68564 | 68553 | 256 | 10 | 4454 | 4466 |
| | NA07357 | 70950 | 70930 | 215 | 18 | 5377 | 5398 |
| | NA07347 | 68676 | 68667 | 244 | 20 | 5427 | 5436 |
| | NA07051 | 68543 | 68533 | 236 | 14 | 4799 | 4810 |
| | NA07048 | 66908 | 66900 | 215 | 13 | 4712 | 4721 |
| | NA06994 | 68066 | 68055 | 234 | 20 | 4959 | 4970 |
| | NA06986 | 70323 | 70308 | 243 | 22 | 4919 | 4934 |
| | NA06984 | 69706 | 69694 | 222 | 20 | 5114 | 5126 |
| $Y_{non\text{-}PARs}$ | NA11831 | 1649 | 1645 | 112 | 0 | 758 | 762 |
| | NA11829 | 3065 | 3051 | 126 | 1 | 1090 | 1104 |
| | NA10851 | 1712 | 1707 | 92 | 2 | 763 | 768 |
| | NA07357 | 1847 | 1838 | 97 | 7 | 822 | 831 |
| | NA07347 | 1684 | 1672 | 124 | 3 | 813 | 825 |
| | NA07051 | 3133 | 3127 | 130 | 5 | 1143 | 1149 |
| | NA07048 | 1816 | 1812 | 104 | 2 | 791 | 795 |
| | NA06994 | 2739 | 2733 | 72 | 2 | 1059 | 1065 |
| | NA06986 | 1946 | 1940 | 112 | 4 | 840 | 846 |
| | NA06984 | 1765 | 1761 | 124 | 4 | 779 | 783 |

### Table S5. Counts of variants across regions of the X and Y.

False negatives are elevated in regions of sequence similarity on the X but evenly dispersed on the Y chromosome**.** Performance metrics for the 10 simulated male and 10 simulated female samples by region.

| **Region** | **Sex** | **Sample** | **# True Positives** | **# False Positives** | **# False Negatives** | **# Simulated** |
|---|---|---|---|---|---|---|
| PARs | Male | NA06984 | 4655 | 1 | 899 | 5554 |
| | | NA06986 | 4932 | 2 | 804 | 5736 |
| | | NA06994 | 5138 | 1 | 808 | 5946 |
| | | NA07048 | 5225 | 6 | 927 | 6152 |
| | | NA07051 | 5032 | 5 | 970 | 6002 |
| | | NA07347 | 5053 | 1 | 919 | 5972 |
| | | NA07357 | 4737 | 3 | 780 | 5517 |
| | | NA10851 | 5058 | 6 | 834 | 5892 |
| | | NA11829 | 4999 | 3 | 938 | 5937 |
| | | NA11831 | 4974 | 1 | 839 | 5813 |
| | Females | NA06985 | 5215 | 3 | 945 | 6161 |
| | | NA06989 | 4759 | 2 | 939 | 5699 |
| | | NA07000 | 5232 | 2 | 828 | 6061 |
| | | NA07037 | 5068 | 2 | 735 | 5803 |
| | | NA07056 | 5026 | 3 | 868 | 5895 |
| | | NA10847 | 4980 | 2 | 886 | 5867 |
| | | NA11830 | 4604 | 1 | 713 | 5318 |
| | | NA11832 | 4632 | 0 | 901 | 5533 |
| | | NA11840 | 5021 | 4 | 944 | 5966 |
| | | NA11892 | 4554 | 5 | 771 | 5326 |
| $X_{\text{non-PARs minus XTR and amplicons}}$ | Males | NA06984 | 66791 | 19 | 4110 | 71531 |
| | | NA06986 | 67696 | 21 | 4068 | 69330 |
| | | NA06994 | 65843 | 19 | 4294 | 70496 |
| | | NA07048 | 63902 | 13 | 3789 | 72487 |
| | | NA07051 | 66170 | 14 | 4022 | 69801 |
| | | NA07347 | 65537 | 18 | 4263 | 70194 |
| | | NA07357 | 68124 | 17 | 4361 | 67695 |
| | | NA10851 | 66513 | 10 | 3981 | 70139 |
| | | NA11829 | 65302 | 13 | 4026 | 71766 |
| | | NA11831 | 67443 | 16 | 4085 | 70903 |
| | Females | NA06985 | 95926 | 136 | 4631 | 100558 |
| | | NA06989 | 97785 | 133 | 5007 | 102793 |

| | | | | | | |
|---|---|---|---|---|---|---|
| | | NA07000 | 99537 | 114 | 5249 | 104787 |
| | | NA07037 | 98161 | 141 | 5176 | 103339 |
| | | NA07056 | 98391 | 116 | 4930 | 103323 |
| | | NA10847 | 96775 | 124 | 4930 | 101707 |
| | | NA11830 | 95508 | 121 | 4722 | 100231 |
| | | NA11832 | 97730 | 112 | 5182 | 102913 |
| | | NA11840 | 98536 | 145 | 4779 | 103316 |
| | | NA11892 | 95575 | 120 | 4975 | 100550 |
| $X_{XTR}$ | Males | NA06984 | 2478 | 1 | 886 | 3364 |
| | | NA06986 | 2241 | 0 | 757 | 2998 |
| | | NA06994 | 1798 | 1 | 567 | 2365 |
| | | NA07048 | 2395 | 0 | 811 | 3206 |
| | | NA07051 | 2016 | 0 | 686 | 2702 |
| | | NA07347 | 2499 | 0 | 1026 | 3525 |
| | | NA07357 | 2422 | 1 | 919 | 3341 |
| | | NA10851 | 1638 | 0 | 386 | 2024 |
| | | NA11829 | 2468 | 2 | 952 | 3420 |
| | | NA11831 | 1754 | 1 | 574 | 2328 |
| | Females | NA06985 | 4411 | 4 | 69 | 4480 |
| | | NA06989 | 4571 | 5 | 95 | 4666 |
| | | NA07000 | 4753 | 8 | 100 | 4853 |
| | | NA07037 | 4856 | 2 | 100 | 4956 |
| | | NA07056 | 4531 | 3 | 98 | 4629 |
| | | NA10847 | 4780 | 2 | 121 | 4901 |
| | | NA11830 | 3980 | 5 | 84 | 4064 |
| | | NA11832 | 4510 | 1 | 86 | 4596 |
| | | NA11840 | 3775 | 2 | 73 | 3848 |
| | | NA11892 | 4984 | 4 | 99 | 5083 |
| $X_{Amplicons}$ | Males | NA06984 | 425 | 0 | 130 | 555 |
| | | NA06986 | 371 | 1 | 109 | 480 |
| | | NA06994 | 414 | 0 | 109 | 523 |
| | | NA07048 | 603 | 0 | 121 | 724 |
| | | NA07051 | 347 | 0 | 102 | 449 |
| | | NA07347 | 631 | 2 | 147 | 778 |
| | | NA07357 | 384 | 0 | 118 | 502 |
| | | NA10851 | 402 | 0 | 99 | 501 |
| | | NA11829 | 274 | 0 | 104 | 378 |
| | | NA11831 | 629 | 1 | 131 | 760 |

| | | | | | | |
|---|---|---|---|---|---|---|
| | Females | NA06985 | 720 | 2 | 195 | 915 |
| | | NA06989 | 793 | 1 | 150 | 943 |
| | | NA07000 | 505 | 2 | 213 | 718 |
| | | NA07037 | 536 | 0 | 178 | 714 |
| | | NA07056 | 767 | 0 | 184 | 951 |
| | | NA10847 | 489 | 0 | 166 | 655 |
| | | NA11830 | 513 | 1 | 151 | 664 |
| | | NA11832 | 526 | 0 | 179 | 705 |
| | | NA11840 | 748 | 1 | 180 | 928 |
| | | NA11892 | 552 | 1 | 170 | 722 |
| $Y_{\text{non-PARs minus XTR and amplicons}}$ | Males | NA06984 | 1607 | 2 | 714 | 2321 |
| | | NA06986 | 1727 | 1 | 763 | 2491 |
| | | NA06994 | 2214 | 2 | 855 | 3070 |
| | | NA07048 | 1559 | 2 | 708 | 2268 |
| | | NA07051 | 2521 | 5 | 928 | 3450 |
| | | NA07347 | 1508 | 3 | 742 | 2251 |
| | | NA07357 | 1666 | 6 | 744 | 2410 |
| | | NA10851 | 1564 | 1 | 686 | 2250 |
| | | NA11829 | 2459 | 0 | 891 | 3351 |
| | | NA11831 | 1462 | 0 | 684 | 2146 |
| $Y_{XTR}$ | Males | NA06984 | 18 | 2 | 13 | 2513 |
| | | NA06986 | 69 | 3 | 17 | 2701 |
| | | NA06994 | 200 | 0 | 81 | 3518 |
| | | NA07048 | 95 | 0 | 18 | 2495 |
| | | NA07051 | 254 | 0 | 82 | 3941 |
| | | NA07347 | 47 | 0 | 22 | 2429 |
| | | NA07357 | 28 | 1 | 16 | 2625 |
| | | NA10851 | 27 | 1 | 16 | 2432 |
| | | NA11829 | 258 | 1 | 83 | 3815 |
| | | NA11831 | 29 | 0 | 12 | 2366 |
| $Y_{ampicons}$ | Males | NA06984 | 136 | 0 | 56 | 192 |
| | | NA06986 | 144 | 0 | 66 | 210 |
| | | NA06994 | 319 | 0 | 129 | 448 |
| | | NA07048 | 158 | 0 | 69 | 227 |
| | | NA07051 | 352 | 0 | 139 | 491 |
| | | NA07347 | 117 | 0 | 61 | 178 |
| | | NA07357 | 144 | 0 | 71 | 215 |
| | | NA10851 | 116 | 0 | 66 | 182 |

| | | | | | | |
|---|---|---|---|---|---|---|
| | | NA11829 | 334 | 0 | 130 | 464 |
| | | NA11831 | 154 | 0 | 66 | 220 |

### Table S6. Using diploid-based filter thresholds for some filters on haploid chromosomes results in less accurate variant calling.

Average number of true positives, false positives, and false negatives on the sex chromosomes for 10 simulated males and 10 simulated females using diploid- and haploid-based thresholds for total depth (DP), allele number (AN), and QualByDepth (QD).

| Filter | Sex | Chromosome | Threshold | Average # TP | Average # FP | Average # FN |
|---|---|---|---|---|---|---|
| DP | Males | $X_{non\text{-}PARs}$ | Diploid based (67 and 201) | 16082 | 2 | 57844 |
| | | | Haploid based (23 and 69) | 58662 | 24 | 15263 |
| | | $Y_{non\text{-}PARs}$ | Diploid based (67 and 201) | 685 | 1 | 2337 |
| | | | Haploid based (23 and 69) | 1944 | 7 | 1103 |
| | Females | $X_{non\text{-}PARs}$ | Diploid based (67 and 201) | 103705 | 210 | 4045 |
| | | | Haploid based (23 and 69) | 771 | 9 | 106689 |
| AN | Males | $X_{non\text{-}PARs}$ | 1 | 71695 | 27 | 2230 |
| | | | 2 | 71692 | 27 | 2233 |
| | | | 3 | 71687 | 27 | 2237 |
| | | | 4 | 71676 | 27 | 2248 |
| | | | 5 | 71655 | 27 | 2270 |
| | | | 10 | 67042 | 23 | 6883 |
| | | | 15 | 0 | 0 | 73927 |
| | | | 20 | 0 | 0 | 73927 |
| | | $Y_{non\text{-}PARs}$ | 1 | 2588 | 10 | 433 |
| | | | 2 | 2586 | 10 | 434 |
| | | | 3 | 2583 | 10 | 437 |
| | | | 4 | 2577 | 10 | 443 |
| | | | 5 | 2569 | 10 | 452 |
| | | | 10 | 2333 | 8 | 688 |
| | | | 15 | 0 | 0 | 3022 |
| | | | 20 | 0 | 0 | 3022 |
| | Females | $X_{non\text{-}PARs}$ | 1 | 104803 | 222 | 2946 |
| | | | 2 | 104803 | 222 | 2946 |
| | | | 3 | 104802 | 222 | 2948 |
| | | | 4 | 104802 | 222 | 2948 |
| | | | 5 | 104800 | 222 | 2950 |

| | | | | | | |
|---|---|---|---|---|---|---|
| | | | 10 | 104788 | 221 | 2962 |
| | | | 15 | 104702 | 220 | 3048 |
| | | | 20 | 104259 | 217 | 3491 |
| QD | Males | $X_{non\text{-}PARs}$ | 1 | 71695 | 27 | 2230 |
| | | | 1.5 | 71695 | 27 | 2230 |
| | | | 2 | 71695 | 27 | 2230 |
| | | | 12 | 71691 | 21 | 2234 |
| | | | 16 | 71684 | 19 | 2241 |
| | | | 20 | 71637 | 17 | 2287 |
| | | | 28 | 54357 | 11 | 19568 |
| | | $Y_{non\text{-}PARs}$ | 1 | 2588 | 10 | 433 |
| | | | 1.5 | 2588 | 10 | 433 |
| | | | 2 | 2586 | 10 | 433 |
| | | | 12 | 2586 | 4 | 435 |
| | | | 16 | 2584 | 3 | 436 |
| | | | 20 | 2578 | 2 | 442 |
| | | | 28 | 1971 | 1 | 1049 |
| | Females | $X_{non\text{-}PARs}$ | 1 | 104803 | 219 | 2946 |
| | | | 1.5 | 104803 | 212 | 2946 |
| | | | 2 | 104801 | 192 | 2949 |
| | | | 12 | 100586 | 13 | 7164 |
| | | | 16 | 90508 | 8 | 17242 |
| | | | 20 | 74247 | 3 | 33503 |
| | | | 28 | 34839 | 1 | 72911 |

### Table S7. Sex chromosome complement specific variants in genes with clinical relevance.

For each gene in which we identified novel variants, we looked in the GWAS Catalog, OMIM phenotypes, and ClinVar to identify whether the gene had previously been implicated in human disease.

| Gene | Var | GWAS Catalog Risk Variant Reported Trait (gene) | OMIM Phenotypes (gene) | ClinVar (gene) |
|---|---|---|---|---|
| PLCXD1 | 139 | - | - | 96 Pathogenic Variants; Conditions: Autism, Schizophrenia, Intellectual Disability |
| GTPBP6 | 114 | - | - | 96 Pathogenic Variants; Conditions: Autism, Schizophrenia, Intellectual Disability |
| PPP2R3B | 1 | - | - | 97 Pathogenic Variants; Conditions: Autism, Schizophrenia, Intellectual Disability |
| SHOX | 195 | Severe COVID-19 infection, Vaginal microbiome | Langer mesomelic dysplasia, Leri-Weill dyschondrosteosis, Short stature, idiopathic familial | 147 Pathogenic Variants; Conditions: Autism, Schizophrenia, Intellectual Disability, SHOX-related short stature, Leri-Weill dyschondrosteosis, Langer mesomelic dysplasia syndrome |
| RPL14P5 | 4 | Schizophrenia, vaginal microbiome | - | - |
| CRLF2 | 185 | Severe COVID-19 infection, Schizophrenia, Eosinophil counts, Eosinophil percentage of white cells | - | 100 Pathogenic Variants; Conditions: Autism, Schizophrenia, Intellectual Disability |

| | | | | |
|---|---|---|---|---|
| CSF2RA | 320 | Mitochondrial heteroplasmy, Granulocyte-macrophage-colony stimulating factor receptor subunit alpha levels, Eosinophil counts, Interleukin-3 receptor subunit alpha levels, Basophil count, Basophil percentage of leukocytes | Surfactant metabolism dysfunction, pulmonary, 4 | 115 Pathogenic Variants; Conditions: Autism, Schizophrenia, Intellectual Disability, Surfactant metabolism dysfunction, pulmonary, 4, CSF2RA-related condition |
| RNA5SP498 | 1 | - | - | - |
| IL3RA | 219 | Interleukin-3 receptor subunit alpha levels, Basophil count, Basophil percentage of leukocytes | - | 96 Pathogenic Variants; Conditions: Autism, Schizophrenia, Intellectual Disability |
| SLC25A6 | 30 | - | - | 95 Pathogenic Variants; Conditions: Autism, Schizophrenia, Intellectual Disability |
| LINC00106 | 36 | - | - | 95 Pathogenic Variants; Conditions: Autism, Schizophrenia, Intellectual Disability |
| ASMTL-AS1 | 46 | - | - | 95 Pathogenic Variants; Conditions: Autism, Schizophrenia, Intellectual Disability |
| ASMTL | 329 | N-acetylserotonin O-methyltransferase-like protein levels | - | 95 Pathogenic Variants; Conditions: Autism, Schizophrenia, Intellectual Disability |
| P2RY8 | 439 | Severe COVID-19 infection, Lymphocyte percentage of leukocytes, Eosinophil counts, Eosinophil percentage of leukocytes, Lymphocyte count, Erythrocyte volume, Mean reticulocyte volume, Erthrocyte volume | - | 95 Pathogenic Variants; Conditions: Autism, Schizophrenia, Intellectual Disability |

| | | | | |
|---|---|---|---|---|
| AKAP17A | 37 | Lymphocyte count | - | 96 Pathogenic Variants; Conditions: Autism, Schizophrenia, Intellectual Disability |
| ASMT | 144 | - | - | 96 Pathogenic Variants; Conditions: Autism, Schizophrenia, Intellectual Disability |
| DHRSX | 1307 | Severe COVID-19 infection, Brain language network functional connectivity | Congenital disorder of glycosylation, type 1DD | 101 Pathogenic Variants; Conditions: Autism, Schizophrenia, Intellectual Disability, Congenital disorder of glycosylation, type 1DD |
| DHRSX-IT1 | 8 | - | - | - |
| ZBED1 | 58 | - | - | 99 Pathogenic Variants; Conditions: Autism, Schizophrenia, Intellectual Disability |
| CD99P1 | 237 | Adolescent idiopathic scoliosis | - | - |
| LINC00102 | 6 | - | - | 98 Pathogenic Variants; Conditions: Autism, Schizophrenia, Intellectual Disability |
| CD99 | 215 | Docking protein 1 levels, erythrocyte volume | - | 98 Pathogenic Variants; Conditions: Autism, Schizophrenia, Intellectual Disability |
| XG | 154 | Glycoprotein Xg levels, Erythrocyte volume | XG blood group system, Xg(a-) phenotype | 208 Pathogenic Variants; Conditions: Autism, Schizophrenia, Kleinfelter's Syndrome, Intellectual disability, Turner's Syndrome, Trisomy X syndrome, 46,XX sex reversal 1, Hypotonia, Neurodevelopmental disorder |
| MXRA5 | 4 | Hip circumference, Severe COVID-19 infection, Intraocular pressure, Pulse pressure | - | 209 Pathogenic Variants; Conditions: Autism, Schizophrenia, Klinefelter syndrome, Turner syndrome, Intellectual Disability, Intellectual Developmental Disorder, Inborn genetic diseases, |
| PRKX | 2 | Thyroid stimulating hormone levels, Blood urea nitrogen levels | - | 204 Pathogenic Variants; Conditions: Autism, Schizophrenia, Klinefelter syndrome, Turner syndrome, Intellectual Disability, Neurodevelopmental disorder, Trisomy X syndrome, 46,XX sex reversal 1, Hypotonia |

| | | | | |
|---|---|---|---|---|
| NLGN4X | 2 | Schizophrenia, Neuroticism, RBC levels of ADP, RBC levels of citrate, Neuroligin-4 X-linked levels, Insomnia, memory decline, circulating fibrinogen levels, Alzheimer's disease, tuberculosis | Autism susceptibility, X-linked 2, Intellectual developmental disorder, X-linked | 204 Pathogenic Variants; Conditions: Autism, Schizophrenia, Klinefelter syndrome, Turner syndrome, Intellectual Disability, Neurodevelopmental disorder, Trisomy X syndrome, 46,XX sex reversal 1, Hypotonia |
| ANOS1 | 18 | Severe COVID-19 infection, Tumor protein p53-inducible protein 13 levels, Serum levels of protein TP53I13, Nipple retraction (>= grade 2) in breast cancer treated with radiotherapy, Facial morphology (factor 17, height of vermillion upper lip) | Hypogonadotropic hypogonadism 1 with or without anosmia (Kallmann syndrome 1) | 250 Pathogenic Variants; Conditions: Autism, Schizophrenia, Intellectual Disability, Hypogonadotropic hypogonadism 1 with or without anosmia, Klinefelter syndrome, Inborn genetic diseases, Turner syndrome, Trisomy X syndrome, 46,XX sex reversal 1, Hypotonia, Neurodevelopmental disorder |
| CTPS2 | 2 | Suicide, Neuroticism | - | 160 Pathogenic Variants; Conditions: Scizophrenia, Autism, Klinefelter syndrome, Turner syndrome, Trisomy X syndrome, 46,XX sex reversal 1, Hypotonia, Neurodevelopmental disorder, Syndromic X-linked intellectual disability Lubs type |

| | | | | |
|---|---|---|---|---|
| DMD | 1 | COVID-19, vaginal microbiome, Body mass index, breast carcinoma, tuberculosis, waist circumference, influenza A, anxiety, gut microbiome, temporomandibular joint disorder, major depressive disorder, educational attainment, blood urea nitrogen, ovarian carcinoma, metabolite levels, migraine, bipolar disorder and schizophrenia, weight, facial morphology, electroencephalogram vigilance, memory performance, itch intensity from mosquito bite, half-life of apixaban, Alzheimer's disease, Type 2 diabetes, systolic blood pressure, sphingosine 1-phosphate RBC levels, mortality, ankle-brachial index, octadecenoic acid RBC levels, ovarian carcinoma, Severe COVID-19 infection, serum levels of protein TENM3, periodontitis | Becker muscular dystrophy, Cardiomyopathy, dilated, 3B, Duchenne muscular dystrophy | 2840 Pathogenic Variants; Conditions: Klinefelter syndrome, Duchenne muscular dystrophy, Becker muscular dystrophy, Polymicrogyria, Congenital adrenal hypoplasia, X-linked, Qualitative or quantitative defects of dystrophin, Abnormality of the musculature, Muscular dystrophy, Myopathy, Cardiovascular phenotype, McLeod neuroacanthocytosis syndrome, Elevated circulating creatine kinase concentration, Dilated cardiomyopathy 3B, Colorectal cancer, DMD-related disorder, Highly elevated creatine kinase, Intermediate muscular dystrophy, Turner syndrome, Ornithine carbamoyltransferase deficiency, Neurodevelopmental disorder, Syndromic X-linked intellectual disability Lubs type, Primary familial hypertrophic cardiomyopathy, Fanconi anemia complementation group A, Becker muscular dystrophy, atypical |

| | | | | |
|---|---|---|---|---|
| KLHL4 | 1 | Severe COVID-19 infection, Bone mineral density, Adolescent idiopathic scoliosis, Alzheimer's disease, Metabolite levels, Alcoholic pancreatitis, memory decline | - | 145 Pathogenic Variants; Conditions: Autism, Schizophrenia, Klinefelter syndrome, Turner syndrome, Trisomy X syndrome, 46,XX sex reversal 1, Hypotonia, Syndromic X-linked intellectual disability Lubs type, Heterotaxy, visceral, 1, X-linked |
| PCDH11X | 349 | Alzheimer's disease, Severe COVID-19 infection, Metabolite levels, Tuberculosis | - | 143 Pathogenic Variants; Conditions: Autism, Schizophrenia, Klinefelter syndrome, Turner syndrome, Intellectual Disability, Neurodevelopmental disorder, Trisomy X syndrome, 46,XX sex reversal 1, Hypotonia, Xq21.32q23 deletion, Developmental and epileptic encephalopathy, 9, Syndromic X-linked intellectual disability Lubs type, Heterotaxy, visceral, 1, X-linked |
| HTR2C | 5 | Educational attainment | - | 154 Pathogenic Variants; Conditions: Autism, Schizophrenia, Klinefelter syndrome, Turner syndrome, Trisomy X syndrome, 46,XX sex reversal 1, Hypotonia, Syndromic X-linked intellectual disability Lubs type, Bone mineral density quantitative trait locus 18, Heterotaxy, visceral, 1, X-linked, Premature ovarian failure |
| SPRY3 | 36 | Body mass index, Triglycerides, Idiopathic osteonecrosis of the femoral head, Total cholestorol levels, Low density liproprotein cholestorol levels, Hemoglobin, Mean corpuscular volume, Mean corpuscular homoglobin, Hematocrit, Red blood cell count, Red cell distribution width, Pyruvate levels in blood donors | - | 104 Pathogenic Variants; Conditions: Autism, Schizophrenia, Kleinfelter's Syndrome, Ectodermal dysplasia and immunodeficiency 1, Immunodeficiency 33, Immunodeficiency 47, Incontinentia pigmenti syndrome, Decreased circulating antibody concentration, Splenomegaly |

| | | | | |
|---|---|---|---|---|
| AMD1P2 | 2 | Pyruvate levels in blood donors | - | - |
| VAMP7 | 126 | Eosinophil counts, Eosinophil percentage of leukocytes | - | 93 Pathogenic Variants; Conditions: Autism, Schizophrenia, Kleinfelter's Syndrome, Ectodermal dysplasia and immunodeficiency 1, Immunodeficiency 33, Immunodeficiency 47, Incontinentia pigmenti syndrome, Decreased circulating antibody concentration, Splenomegaly |
| ELOCP24 | 1 | Eosinophil counts, Eosinophil percentage of leukocytes | - | - |
| TRPC6P | 10 | - | - | - |
| IL9R | 26 | Eosinophil counts, Eosinophil percentage of leukocytes | - | 81 Pathogenic Variants; Conditions: Autism, Schizophrenia, Klinefelter syndrome, Immunodeficiency 47, Immunodeficiency 33, Ectodermal dysplasia and immunodeficiency 1, Incontinentia pigmenti syndrome, Decreased circulating antibody concentration, Splenomegaly, Autoimmune thrombocytopenia |
| AL158055 .1 | 1 | - | - | - |
| AC073614 .1 | 1 | - | - | - |
| AL121872 .1 | 22 | - | - | - |
| AL683807 .1 | 165 | - | - | - |
| AL672277 .1 | 18 | - | - | - |
| AL954722 .1 | 16 | - | - | - |
| AL683807 .2 | 12 | - | - | - |

### Supplementary Figures

#### Figure S1. Variant calling accuracy declines substantially in XTR when masking XTR on the Y chromosome for alignment.

Box plots of the number of called variants over the number of simulated variants in XTR across 10 simulated male (XY) individuals using different alignment approaches. We aligned simulated reads from 10 male (XY) samples to two different versions of the reference genome - one where the PARs on the Y chromosome are hard masked (SCC), and another with both PARs and XTR hard masked on the Y chromosome (SCC + Y-XTR masked) - and called variants and hard filtered using GATKs best practices protocol. We called variants in XTR for the SCC aligned data as haploid for both the X and Y chromosomes and for the SCC + Y-XTR masked aligned data we called variants in XTR as diploid on chromosome X. We then counted the number of variants called in XTR and compared that to the number of simulated variants in XTR for X and Y for each simulated sample. Since variants were called only on X in the approach with XTR hard masked on Y, we summed the number of X and Y chromosome simulated variants.

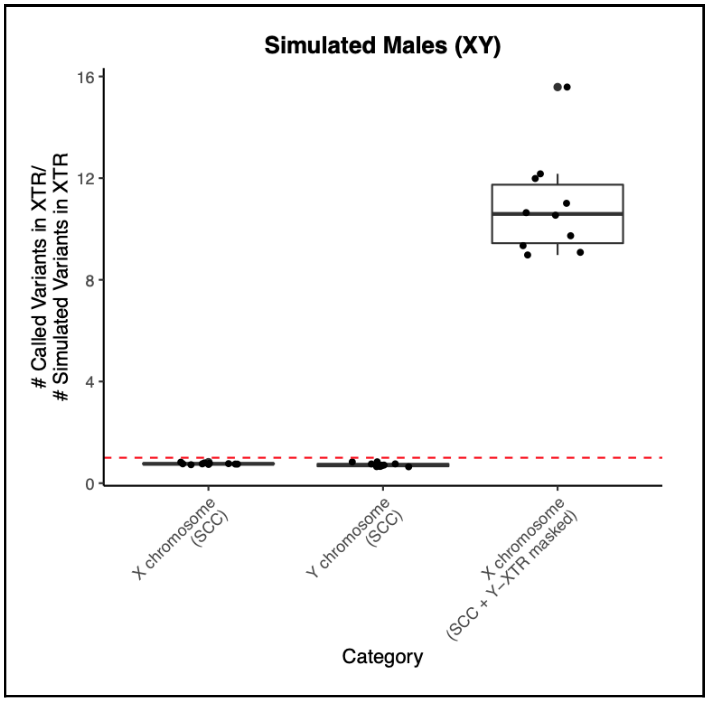

### Figure S2. Proportion of false positives and false negatives on chromosome X and chromosome Y.

The proportion of false positives and false negatives on chromosome X are similar to autosomal ranges while chromosome Y has an elevated proportion of false negatives. Box plots of the proportion of false positives to the total number of simulated variants across A) 10 simulated female (XX) individuals and B) 10 simulated male (XY) individuals, and the proportion of false negatives to the total number of simulated variants across C) 10 simulated female (XX) individuals and D) 10 simulated male (XY) individuals across the autosomes and sex chromosomes. All data was aligned to a reference genome informed on the sex chromosome complement of the sample, diploid calling mode was performed on the autosomes and the X chromosome in simulated females, and haploid calling mode was performed on the X and Y chromosomes in simulated males. GATK hard filters with recommended thresholds were implemented for all chromosomes. Joint genotyping was performed across 10 simulated females and 10 simulated males separately. Highest and lowest proportions of false positives and false negatives are annotated with dashed horizontal red lines.

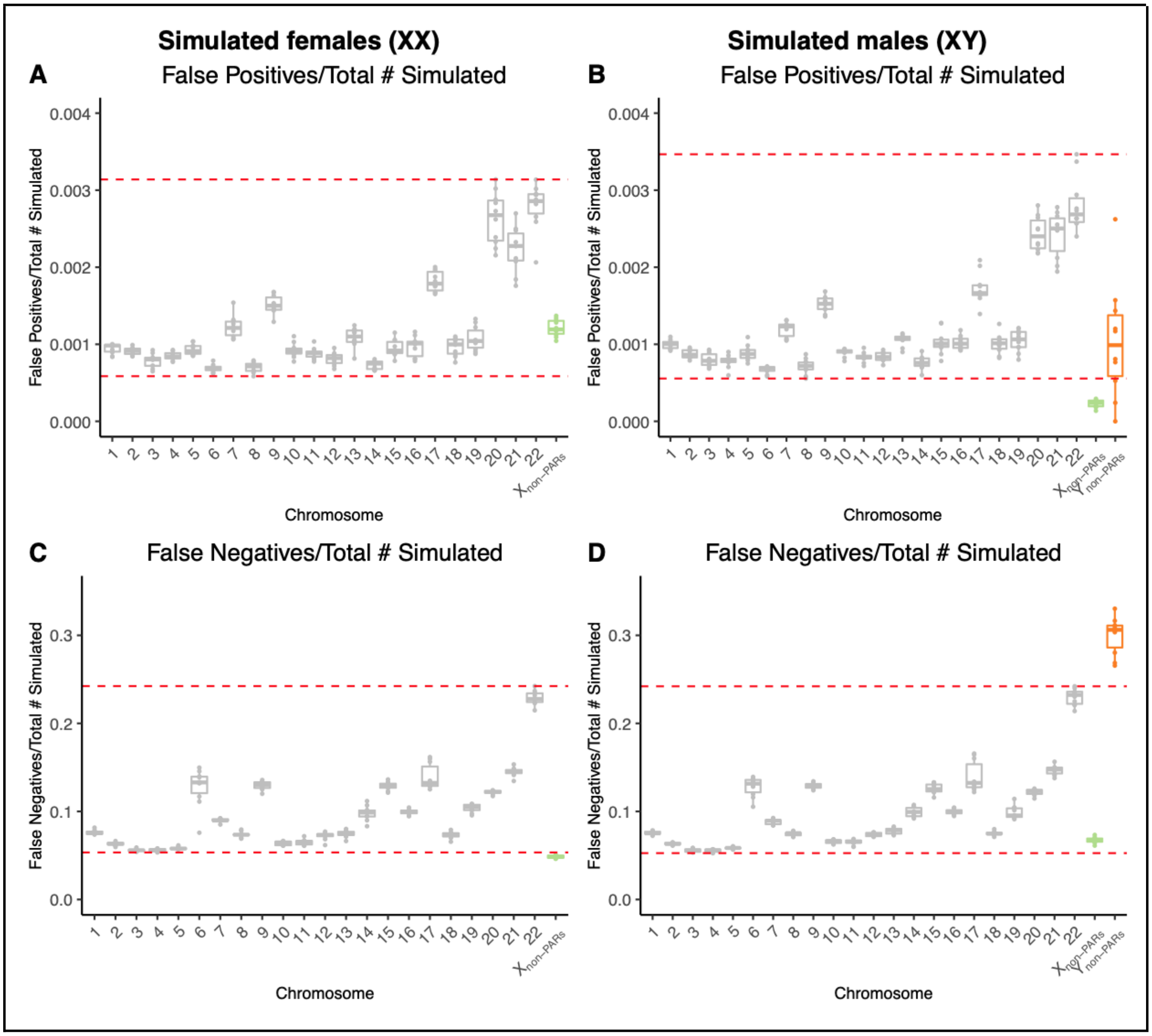

### Figure S3. Distribution of DP, AN, and QD are similar between autosomes and the X chromosome in females

Distributions of variants (top) and the number of true positives and false positives for different filtering thresholds for autosomes, X non-PARs, and Y non-PARs across simulated female (XX) samples for A) total depth, B) allele number, and C) QualByDepth. Chromosome 8 was used to represent the autosomes. Vertical red dashed lines represent filter thresholds based on the autosomes.

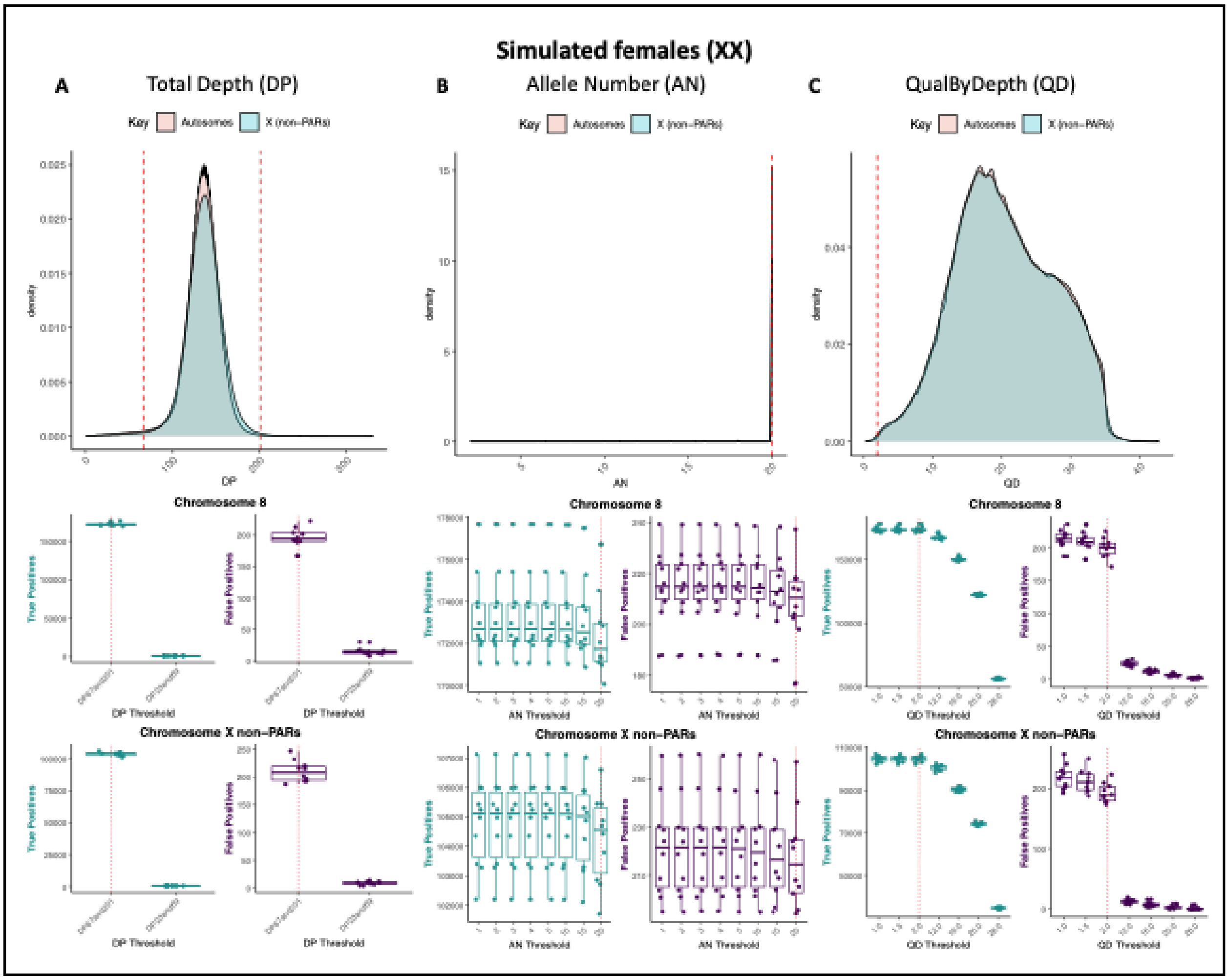

### Figure S4. Comparing 10 vs 20 sample results.

The proportion of false positives to the total number of simulated variants and false negative relative to the total number of simulated variants are similar when joint genotyping across 10 and 20 samples. Box plots of the proportion of false positives to the total number of simulated variants across A) simulated females (XX) and B) simulated males (XY), and the proportion of false negatives to the total number of simulated variants across C) simulated females (XX), D) simulated males (XY) when joint genotyping 10 and 20 samples.

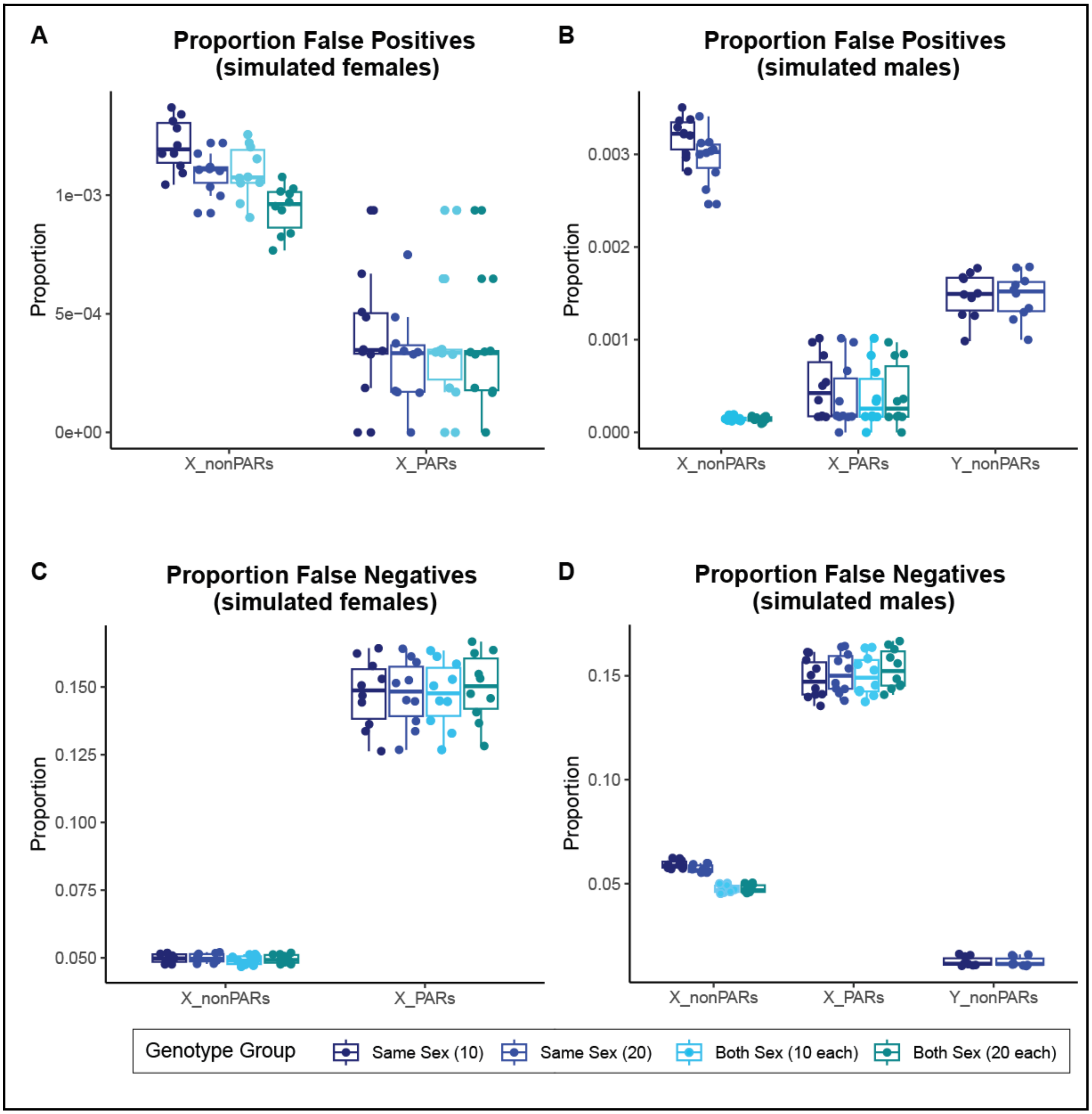

### Figure S5. Proportions of false positives and false negatives are similar across simulated individuals from different ancestries.

Overall, we observed no large differences in the proportion of false positives and false negatives across simulated individuals from different ancestries. In females the proportion of false positives on the PARs are the highest for simulated African samples (average proportion of false positives = 0.0007) but slightly lower for the simulated Asian (average proportion of false positives = 0.0005) and European samples (average proportion of false positives = 0.0004; **Figure S5A**). A similar pattern was observed in males, where the average proportion of false positives is 0.0006 across simulated African and Asians, and 0.0004 across simulated Europeans samples (**Figure S5B**). The proportion of false positives on X non-PARs in females is equal across simulated Asian and European samples (average proportion of false positives = 0.0011) but slightly lower across the simulated African samples (average proportion of false positives = 0.0008; **Figure S5A**). For males, the average proportion of false positives on X non-PARs is equal across all simulated ancestries (average proportion of false positives = 0.0002). On the Y non-PARs, the proportion of false positives is highest for simulated African samples (average proportion of false positives = 0.0011) but slightly lower and equal for the simulated Asian and European samples (average proportion of false positives = 0.0008; **Figure S5B**).

Similar to false positives, the proportion of false negatives on the PARs are highest for simulated African samples (average proportion of false positives = 0.1661) but slightly lower for the simulated Asian (average proportion of false positives = 0.1627) and European female samples (average proportion of false positives = 0.1502; **Figure S5C**). This pattern was also observed across males on the PARs (average proportion of false positives Africans = 0.1644; average proportion of false positives Asians = 0.1622; average proportion of false positives Europeans = 0.1500; **Figure S5D**). On X non-PARs, the proportion of false negatives is lowest across simulated African samples and slightly higher for both Asian and European samples for both simulated females (average proportion of false positives in simulated Africans = 0.0445; average proportion of false positives in simulated Asians = 0.0513; average proportion of false positives in simulated Europeans = 0.0490) and males (average proportion of false positives in simulated Africans = 0.0613; average proportion of false positives in simulated Asians = 0.0692; average proportion of false positives in simulated Europeans = 0.0652; **Figure S5D**). A similar pattern was observed on the Y non-PARs (average proportion of false positives in simulated Africans = 0.2678; average proportion of false positives in simulated Asians = 0.2704; average proportion of false positives in simulated Europeans = 0.2881; **Figure S5D**).

Box plots of the proportion of false positives to the total number of simulated variants across A) 20 simulated females (XX) and B) 20 simulated males (XY), and the proportion of false negatives to the total number of simulated variants across C) 20 simulated females (XX) and D) 20 simulated males (XY) for simulated African, Asian, and European samples.

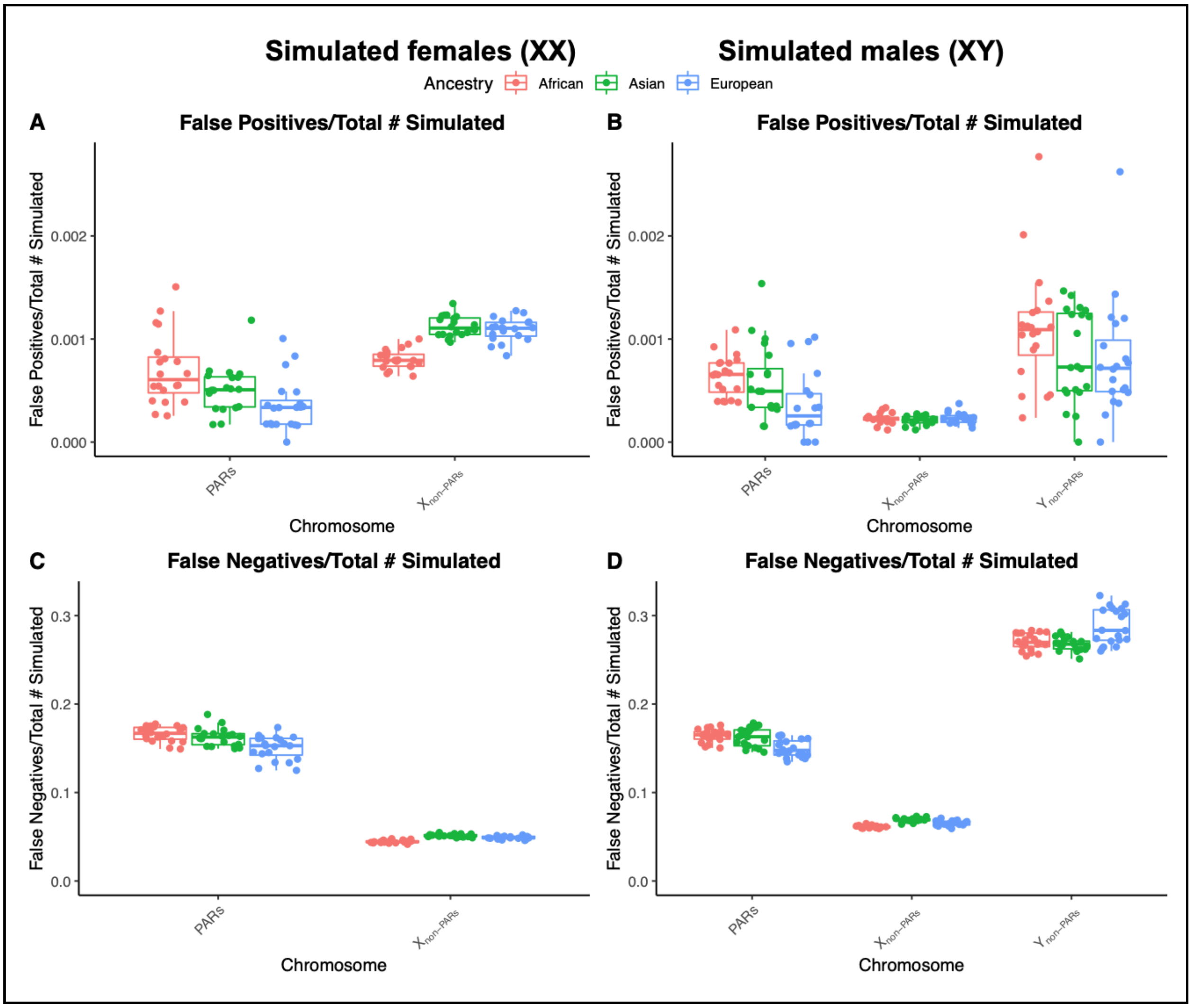

Simulated females (XX)
Simulated males (XY)
Ancestry
African
Asian
European
A
False Positives/Total # Simulated
B
False Positives/Total # Simulated
False Positives/Total # Simulated
0.002
0.001
0.000
PARs
Xnon-PARs
Ynon-PARs
Chromosome
C
False Negatives/Total # Simulated
D
False Negatives/Total # Simulated
False Negatives/Total # Simulated
0.3
0.2
0.1
0.0
